# Same same but different: Cluster architecture variation in five ‘Pinot Noir’ clonal selection lines correlates with differential expression of three transcription factors and further growth related genes

**DOI:** 10.1101/2020.03.17.993907

**Authors:** Robert Richter, Susanne Rossmann, Doreen Gabriel, Reinhard Töpfer, Klaus Theres, Eva Zyprian

## Abstract

Grapevine (*Vitis vinifera* L.) is an economically important crop that needs to comply with high quality standards for fruit, juice and wine production. Intense plant protection is required to avoid losses caused by fungal infections. Grapevine cultivars with loose cluster architecture enable to reduce protective chemical treatments due to their enhanced resilience against fungal infections such as *Botrytis cinerea* induced grey mold. A recent study identified transcription factor gene *VvGRF4* as determinant of inflorescence structure in exemplary samples of loose and compact quasi-isogenic ‘Pinot Noir’ clones. Here, we extended the analysis to 12 differently clustered ‘Pinot Noir’ clones originating from five different clonal selection programs. Differential gene expression of these clones was studied in three different locations over three seasons in demonstrative vineyards. Two phenotypically contrasting clones were grown at all three locations and served for standardization of downstream analyses. Differential gene expression data were correlated to the phenotypic variation of cluster architecture sub-traits. A consistent differential gene expression of *VvGRF4* in relation to loose clusters was verified over the different environments and in the extended set of ‘Pinot Noir’ clones. In addition, 14 more genes with consistent expression differences between loosely and compactly clustered clones independent from season and location were identified. These genes show annotations related to cellular growth, cell wall extension, cell division and auxin metabolism. They include two more transcription factor genes.

## Introduction

Grapevine (*Vitis vinifera* L.) is one of the most important fruit crops at global scale. The worldwide grape production reached 75.8 million tons in 2016 (OIV 2017a). The world gross production value for grapes was above 67.5 billion USD (FAOSTAT 2016). Regardless of the use as wine grapes, table grapes, or dried fruits (raisins) only high quality fruits are acceptable for marketing. Unfortunately, *V. vinifera* grapevine varieties are susceptible to several pathogens and viticulture requires intense application of plant protection products (PPP) to meet the market’s requirements. Fungicides are unavoidable to control the pathogens (Pertot et al. 2017) causing powdery mildew, *Erysiphe necator* (syn. *Uncinula necator*, (Schw.) Burr), downy mildew, *Plasmopara viticola* (Berk. & Curt) Berl. & de Toni) and *Botrytis cinerea* (teleomorph *Botryotinia fuckeliana* ((de Bary) Whetzel), provoking grey mold. The use of PPP, irrespective of their inorganic (copper and sulfur) or synthetic origin, contributes to a decrease in biodiversity and raises consumers concerns (Keulemans et al. 2019). One strategy to reduce their use is the breeding of pathogen-resistant grapevine varieties, e.g. by introgression of genetically seizable resistance loci against *Erysiphe necator* and *Plasmopara viticola* from wild *Vitis spec.* relatives into *Vitis vinifera* quality-cultivars. In the last years, several improved varieties with resistance traits against the mildews became available. However, for *Botrytis cinerea*, there is only preliminary knowledge on a putative resistance locus (Sapkota et al. 2019). Current cultivar development therefore focuses on the enforcement of physical barriers, e.g. a thick berry skin, a hydrophobic berry surface and loose cluster architecture, to increase resilience towards *B. cinerea* (Gabler et al. 2003; Herzog et al. 2015; Shavrukov et al. 2004). Within a loose grape cluster, improved ventilation accelerates the drying-off after rainfall or morning dew. Reduced humidity diminishes infections with fungal pathogens (Hed et al. 2009; Molitor et al. 2012). In addition, fungicide sprays can better spread into a loosely clustered bunch as compared to a compact one (Hed et al. 2010). The high physical stress arising in between the berries of compact clusters upon ripening provokes micro cracks or even bursting of the berry skin (Becker and Knoche 2012; Smart and Robinson 1991). This problem is avoided in loosely clustered bunches. Moreover, there are less pronounced temperature gradients within loosely structured clusters as solar radiation can better reach the interior berries. This conveys more uniform fruit maturity (Pieri et al. 2016; Vail and Marois 1991). Overall, loose cluster architecture results in grapes with less *Botrytis cinerea* infections and a better harmonized ripening process. It is a highly desired trait in grapevine breeding. Understanding its genetic basis should help to develop novel tools for efficient grapevine breeding and clonal selection.

Worldwide, several thousands of grapevine cultivars exist. They are registered in data repositories, e.g. the “*Vitis* International Variety Catalogue” (http://www.vivc.de) (Maul 2019). The gene pools of wine grapes and table grapes show remarkable differences in berry- and cluster architecture (Di Genova et al. 2014; Migicovsky et al. 2017). Despite this impressive genetic diversity, only 33 (*Vitis vinifera* L. subsp. *vinifera*) cultivars account for 50% of the totally used acreage for commercial production (OIV 2017). Within these predominant cultivars, intra-varietal genetic variation, caused by somatic mutation (De Lorenzis et al. 2017), is exploitable to select clonal variants of the desired cluster architecture phenotype.

Bunch architecture is controlled by environmental and genetic factors (Döring et al. 2015; Tello and Ibáñez 2017). It is a complex trait determined by the interplay of berry- and stalk characteristics (Li et al. 2019; Richter et al. 2018; Rist et al. 2018). Some of these sub-traits are under genetic control as reported for berry size, berry volume and berry weight (Ban et al. 2016; Houel et al. 2015; Mejia et al. 2007; Tello et al. 2015), berry number (Dry et al. 2010; Fanizza et al. 2005) and other rachis sub-traits (Correa et al. 2014; Marguerit et al. 2009; Tello et al. 2016).

Intravarietal diversity in cluster architecture sub-traits of grapevine cultivars has been reported in only a few cases, like ‘Garnacha Tinta’, ‘Tempranillo’, ‘Aglianico’ and ‘Muscat of Alexandria’ (Grimplet et al. 2019; Grimplet et al. 2017). For ‘Albariño’ clones and for ‘Pinot Noir’ clones the studies of Alonso-Villaverde et al. (2008) and Konrad et al. (2003) proved, that intravarietal cluster architecture variance correlates with the suseptebility to *B.cinerea*, i.e. loosely clusterd clones show reduced susceptibility. In ‘Pinot Noir’ (PN), the gene *VvGRF4* was recently detected as a major component affecting inflorescence architecture (Rossmann et al. 2019). PN is a member of the very old ‘Pinot’ family (Regner et al. 2000) and is used in viticulture since centuries. Presently, with an area of 115.000 ha, PN is among the top thirteen international varieties (OIV 2017). The ‘Pinot’ family accumulated a high number of somatic mutations and gave rise to a wide range of clones displaying divergent phenotypic features e.g. different berry color, varying organoleptic appearance, different vigor and cluster architecture (Forneck et al. 2009). Concerning cluster architecture (CA), the clones were classified into three categories, i.e. compact (CCC), loose (LCC) and mixed berry type (MBC) ‘Pinot Noir’ clones (Bleyer 2001; Ruehl et al. 2004).

In the previous study, two loosely clustered PN clones from the “Mariafeld” selection line (M171) and the Geisenheim clonal selection program (Gm1-86) were compared to two compactly clustered clones (“Frank Charisma” and “Frank Classic”). This investigation revealed a mutation in the micro RNA mi396 binding site of *VvGRF4*, a gene encoding a growth promoting transcription factor. The mutation prevents down-regulation of the *VvGRF4* transcript specifically in the LCC clones. Two mutated alleles were identified, one specific for M171, the other one found in Gm1-86. Both operate in heterozygous state, lead to an enhancement of cell numbers in pedicels in the loose clusters and thus contribute to loose cluster architecture (Rossmann et al. 2019). This study here explored variation of cluster architecture in combination with differential transcriptional activity in an extended set of twelve PN clones from different selection lines. Besides other genes, the activity of *VvGRF4* was investigated to check its relevance in further PN clones and over several environments.

A detailed morphological characterization was undertaken ahead to reveal the relevant sub-traits of cluster architecture within the PN clonal groups. These sub-traits were indexed to group the clones into loose and compact clones. Expression levels of genes selected from a previous RNA-Seq study including *VvGRF4* (Rossmann et al. 2019) or literature references were then interrogated to identify differentially expressed genes involved in the expression of bunch compactness. Broadening the previous work, environmental effects were taken into account. PN plants from three geographically distinct trial plots, managed with organic and integrated practices, were investigated over three consecutive growing seasons. In addition to *VvGRF4*, this investigation revealed additional genes involved in the determination of cluster architecture, acting independently from environmental factors (season, location, vineyard management) in an extended and diverse clonal set of PN. These newly identified genes encode two more transcription factors and functions related to auxin metabolism and cellular growth.

## Material and Methods

### Plant material

The *Vitis vinifera* variety ‘Pinot Noir’ was investigated in 12 clones showing divergent cluster architecture. These comprised compactly clustered clones (CCCs), loosely clustered clones (LCCs) and clones bearing berries with mixed size (MBCs). The plants were distributed over three plantations in three German viticulture areas (Palatinate, Baden and Hesse) with partial overlapping redundancy (Table 1). The vineyards in Baden and Hesse were managed by grapevine nurseries and originated from certified material. They were submitted to regular visual monitoring for their phytosanitary state. The PN clones were well established (∼20 year old vines) and all grafted on the same rootstock (Kober 125AA). “Guyot pruning” was applied throughout and a vertical shoot position trellis system with 1.8 to 2.2m^2^ space per vine was used. Vineyards in Baden and Hesse were maintained with integrated management. The Palatinate trial field belongs to the Julius Kuehn Institute for Grapevine Breeding Geilweilerhof. This one was managed according to organic farming rules (Döring et al. 2015) (Online resource 1). All the plantations contained ample material of individual PN plants to permit random sampling from the individual clones. Samples were taken exclusively from plants without any symptom of infection or aberration from the typical clonal type of appearance.

**Table 1.** Sampling schedules for 12 ‘Pinot Noir’ clones spread over three locations during three seasons. For phenotyping of cluster traits, samples of ripe bunches at BBCH89 were taken with 10 replicates from randomly selected independent vines. The measurements of the PN clones ‘Frank Charisma’ (FkCH) and ‘Gm20-13’, present at all three locations, enabled to model the environmental impact on cluster architecture sub-traits (Online resource 3 a and b).

|  |  |  | Palatinate | Hesse | Baden |
| --- | --- | --- | --- | --- | --- |
| Cluster type | ‘Pinot Noir’ clone | abbreviation | BBCH 89 | BBCH 89 | BBCH 89 |
| CCC | **Frank Charisma** | FkCH | 10<sup>a</sup> | 10<sup>a</sup> | 10<sup>a</sup> |
| CCC | Frank Classic | FkCL | 10<sup>a</sup> | 10<sup>a</sup> | - |
| CCC | Entav 777 | En777 | - | 10<sup>a</sup> | 10<sup>a</sup> |
| variable | Geisenheim 18 | Gm18 | - | 10<sup>b</sup> | - |
| MBC | **Geisenheim 20-13** | Gm20-13 | 10<sup>a</sup> | 10<sup>a</sup> | 10<sup>a</sup> |
| MBC | Freiburg 1801 | Fr1801 | - | 10<sup>a</sup> | 10<sup>b</sup> |
| LCC | Geisenheim 1-86 | Gm1-86 | 10<sup>a</sup> | 10<sup>a</sup> | - |
| LCC | Freiburg 12-L | Fr12L | - | 10<sup>a</sup> | 10<sup>a</sup> |
| LCC | Freiburg 13-L | Fr13L | - | 10<sup>a</sup> | 10<sup>a</sup> |
| LCC | Weinsberg M1 | WeM1 | - | 10<sup>a</sup> | - |
| LCC | Weinsberg M171 | WeM171 | 10<sup>a</sup> | - | - |
| LCC | Weinsberg M242 | WeM242 | - | 10<sup>b</sup> | - |
| a= ten biological samples taken in 2015 and 2016 |  |  |  |  |  |
| b= ten biological samples taken in 2016 |  |  |  |  |  |
| -= not available |  |  |  |  |  |

### Sampling

For the phenotypic evaluation at BBCH89 (berries ripe for harvest), ten vines per clone were chosen randomly. From every vine, a basally inserted cluster from the central shoot of the fruit cane was collected in the years 2015 and 2016 at each vineyard. Bunches were cut directly at the connection with the shoot and stored at 5°C until use. Samples for gene expression experiments were taken in the same way (Table 2), but collected as triplicates at the early developmental stages BBCH57 (just before flowering) and BBCH71 (at early fruit set) during the three years from 2015 to 2017. Complete inflorescences were cut and shock-frozen immediately in liquid nitrogen. The non-linear cumulative degree-day (CDD) based model (Molitor et al. 2014) adjusted the sampling to ensure the same developmental stage over the three locations studied during all three years. The target temperature sum was 400° CDD for BBCH57 and 700° CDD for BBCH71. CDD calculation was based on air temperatures at 2m height recorded by the nearest weather station. A detailed schedule of the sampling and the temperature records is presented in online resource 2.

**Table 2.** Sampling schedule for differential gene expression analysis

|  |  |  | Palatinate |  | Hesse |  | Baden |  |
| --- | --- | --- | --- | --- | --- | --- | --- | --- |
|  |  |  | BBCH |  | BBCH |  | BBCH |  |
| Cluster type | 'Pinot Noir' clone | abbreviation | 57 | 71 | 57 | 71 | 57 | 71 |
| CCC | **Frank Charisma** | FkCH | 3<sup>a</sup> | 3<sup>a</sup> | 3<sup>a</sup> | 3<sup>a</sup> | 3<sup>a</sup> | 3<sup>a</sup> |
| CCC | Frank Classic | FkCL | 3<sup>a</sup> | 3<sup>a</sup> | 3<sup>a</sup> | 3<sup>a</sup> | - | - |
| CCC | Entav 777 | En777 | - | - | 3<sup>a</sup> | 3<sup>a</sup> | 3<sup>a</sup> | 3<sup>a</sup> |
| unsteady | Geisenheim 18 | Gm18 | - | - | 3<sup>b</sup> | 3<sup>b</sup> | - | - |
| **MBC** | **Geisenheim 20-13** | Gm20-13 | 3<sup>a</sup> | 3<sup>a</sup> | 3<sup>a</sup> | 3<sup>a</sup> | 3<sup>a</sup> | 3<sup>a</sup> |
| MBC | Freiburg 1801 | Fr1801 | - | - | 3<sup>b</sup> | 3<sup>b</sup> | 3<sup>a</sup> | 3<sup>a</sup> |
| LCC | Geisenheim 1-86 | Gm1-86 | 3<sup>a</sup> | 3<sup>a</sup> | 3<sup>a</sup> | 3<sup>a</sup> | - | - |
| LCC | Freiburg 12-L | Fr12L | - | - | 3<sup>b</sup> | 3<sup>b</sup> | 3<sup>b</sup> | 3<sup>b</sup> |
| LCC | Freiburg 13-L | Fr13L | - | - | 3<sup>b</sup> | 3<sup>b</sup> | 3<sup>b</sup> | 3<sup>b</sup> |
| LCC | Weinsberg M1 | WeM1 | - | - | 3<sup>b</sup> | 3<sup>b</sup> | - | - |
| LCC | Weinsberg M171 | WeM171 | 3<sup>a</sup> | 3<sup>a</sup> | - | - | - | - |
| LCC | Weinsberg M242 | WeM242 | - | - | 3<sup>b</sup> | 3<sup>b</sup> | - | - |
| a= three biological samples taken in 2015, 2016 and 2017<br>b= three biological samples taken in 2016 and 2017<br>-= not available |  |  |  |  |  |  |  |  |

### Evaluation of vegetative growth

Vitality of the PN clones was assessed by measuring the mass of the annual outgrowth i.e. the weight of the ten most basally located branches on ten vines per season and location (Online resource 1, Table 3).

**Table 3.** Morphometric measurements on cluster architecture for 12 ‘Pinot Noir’ clones at BBCH89 recorded over three locations and two seasons. Estimated (marginal) means of sub-traits and compactness indices for each clone adjusted for the effects of “location” and “season” as predicted from the generalized linear model “sub-trait” ∼ loc*year+clone (details in Online resource 2). (±) represents the standard error. Different letters indicate significantly divergent values for sub-traits and compactness indices as identified with a Tukey HSD test at significance level α = 0.05.

| Trait | Abbreviation | En777 | FkcH | FkcL | Fr12L | Fr13L | Fr1801 | Gm20-13 | Gm1-86 | WeM1 | WeM171 | WeM242 | Gm18 |
| --- | --- | --- | --- | --- | --- | --- | --- | --- | --- | --- | --- | --- | --- |
|  |  | compact | compact | compact | loose | loose | loose | loose | loose | loose | loose | loose | unsteady |
| Peduncle length | PL [cm] | 1.24<br>(±0.1)<br>abcd | 1.16<br>(±0.07)<br>abc | 1.13<br>(±0.09)<br>ab | 1.38<br>(±0.11)<br>abcd | 1.58<br>(±0.11)<br>bcd | 1.14<br>(±0.11)<br>abc | 1.02<br>(±0.08)<br>a | 1.72<br>(±0.12)<br>d | 1.65<br>(±0.17)<br>bcd | 1.42<br>(±0.16)<br>abcd | 1.93<br>(±0.27)<br>cd | 1.05<br>(±0.19)<br>abcd |
| Rachis weight | RW [g] | 7.4<br>(±0.36)<br>abc | 8.76<br>(±0.28)<br>cde | 8.25<br>(±0.36)<br>abcd | 8.94<br>(±0.37)<br>cde | 8.21<br>(±0.36)<br>abcd | 8.12<br>(±0.42)<br>abcd | 6.69<br>(±0.31)<br>a | 9.62<br>(±0.36)<br>de | 6.78<br>(±0.52)<br>ab | 8.72<br>(±0.53)<br>bcde | 10.97<br>(±0.73)<br>e | 7.81<br>(±0.73)<br>abcde |
| Rachis diameter | RD[cm] | 0.4<br>(±0.01)<br>bc | 0.35<br>(±0.01)<br>a | 0.39<br>(±0.01)<br>bc | 0.4<br>(±0.01)<br>bc | 0.4<br>(±0.01)<br>bc | 0.38<br>(±0.01)<br>abc | 0.39<br>(±0.01)<br>b | 0.42<br>(±0.01)<br>c | 0.38<br>(±0.01)<br>ab | 0.38<br>(±0.01)<br>abc | 0.43<br>(±0.02)<br>bc | 0.37<br>(±0.02)<br>abc |
| first internode length | L1I [cm] | 1.27<br>(±0.11)<br>a | 1.31<br>(±0.09)<br>a | 1.26<br>(±0.11)<br>a | 1.53<br>(±0.12)<br>a | 1.26<br>(±0.11)<br>a | 1.6<br>(±0.13)<br>a | 1.3<br>(±0.1)<br>a | 1.45<br>(±0.11)<br>a | 1.53<br>(±0.16)<br>a | 1.46<br>(±0.17)<br>a | 2.08<br>(±0.23)<br>a | 1.45<br>(±0.23)<br>a |
| second internode length | L2I [cm] | 1.28<br>(±0.09)<br>a | 1.27<br>(±0.07)<br>a | 1.29<br>(±0.09)<br>a | 1.49<br>(±0.09)<br>a | 1.54<br>(±0.09)<br>a | 1.13<br>(±0.11)<br>a | 1.21<br>(±0.08)<br>a | 1.37<br>(±0.09)<br>a | 1.46<br>(±0.13)<br>a | 1.47<br>(±0.14)<br>a | 1.49<br>(±0.19)<br>a | 1.35<br>(±0.19)<br>a |
| seasonal wood gain | WG [g] | 790<br>(±30)<br>bc | 716<br>(±21)<br>abc | 613<br>(±23)<br>a | 702<br>(±27)<br>abc | 672<br>(±25)<br>ab | 807<br>(±36)<br>bc | 790<br>(±26)<br>bc | 807<br>(±31)<br>c | 677<br>(±37)<br>abc | 676<br>(±38)<br>abc | 693<br>(±53)<br>abc | 755<br>(±58)<br>abc |
| <sup>a</sup>Index | BN/RL [cm] | 14.39<br>(±0.51)<br>f | 12.73<br>(±0.35)<br>ef | 12.81<br>(±0.45)<br>ef | 10.57<br>(±0.38)<br>bcd | 10.77<br>(±0.37)<br>bcd | 8.84<br>(±0.36)<br>a | 10.94<br>(±0.33)<br>bcd | 12.01<br>(±0.42)<br>de | 9.8<br>(±0.49)<br>abc | 9.48<br>(±0.49)<br>ab | 10.16<br>(±0.72)<br>abcde | 12.71<br>(±0.7)<br>cdef |
| <sup>b</sup>Index | **CI-12** | 1.49<br>(±0.07)<br>**f** | 1.19<br>(±0.04)<br>**ef** | 1.17<br>(±0.05)<br>**de** | 0.99<br>(±0.05)<br>**cde** | 1.04<br>(±0.05)<br>**cde** | 0.63<br>(±0.03)<br>**a** | 0.78<br>(±0.03)<br>**b** | 1.17<br>(±0.05)<br>**de** | 0.93<br>(±0.06)<br>**bcd** | 0.87<br>(±0.06)<br>**bc** | 0.87<br>(±0.08)<br>**abcd** | 1.06<br>(±0.09)<br>**bcde** |
| <sup>c</sup>Index | **CI-18** | 4.91<br>(±0.43)<br>**f** | 3.24<br>(±0.22)<br>**ef** | 3.24<br>(±0.28)<br>**de** | 1.77<br>(±0.16)<br>**abc** | 1.96<br>(±0.17)<br>**abc** | 1.31<br>(±0.13)<br>**a** | 2.26<br>(±0.17)<br>**bcd** | 2.31<br>(±0.2)<br>**bcde** | 1.58<br>(±0.2)<br>**ab** | 1.89<br>(±0.24)<br>**abc** | 1.33<br>(±0.23)<br>**ab** | 3.26<br>(±0.57)<br>**cdef** |

<sup>a</sup> According to Hed et al. (2009) <sup>b</sup> According to Tello and Ibáñez (2014) <sup>c</sup> based on CI-18 stated in Tello and Ibáñez (2014) but omitting seed number. cluster architecture sub-traits indicated in bold are major contributors to cluster density levels (Richter et al. 2018)

### Phenotypic evaluation of cluster architecture sub-traits

Measurements of 12 cluster architecture sub-traits (Table 3) evaluated the phenotypes. Three indices for cluster compactness were calculated. The ratio “berry number/rachis length” (BN/RL[cm], Hed et al., 2009), and indices CI-12 (berry weight [g] / (rachis length [cm])^2^ and CI-18 (berry weigth [g] x berry number/ (peduncle length [cm] + rachis length [cm])^2^ x rachis length [cm] x pedicel length [mm]) followed the suggestion of Tello and Ibáñez (2014).

### RNA extraction and cDNA synthesis

The pre-bloom flowers (BBCH57), respectively fruit setting berries (BBCH71), were carefully removed from the samples. The complete remaining rachis structure was ground into fine powder. All steps were performed in liquid nitrogen. Aliquots of sample tissue were mixed with 500 mg polyvinylpyrrolidone Polyclar® AT (Serva Electrophoresis GmbH, Heidelberg, Germany). Total RNA extraction used the Spectrum™ Plant Total RNA Kit (Sigma Aldrich, Darmstadt, Germany), following protocol “A”. An on-column DNaseI digestion with RNase-Free DNase (QIAGEN, Hilden, Germany), was performed according to the manufacturer’s protocol. RNA integrity and quantity were analyzed by spectrophotometry (Clario Star 0430, BMG Labtech, Ortenberg, Germany) and checking 500ng of total RNA by non-denaturing agarose gel (1%) electrophoresis. 250ng of total RNA was used for first-strand cDNA synthesis with the High capacity cDNA Transcription Kit (Applied Biosystems, ThermoFisher Scientific, Waltham, MA, USA) following the manufacturer’s protocol.

### Primer design

Primers (listed in Online resource 3) were designed as recommended in (Citri et al. 2012) using the CLC-main workbench primer design software tool (CLC Main Workbench Version 8.0.1, QIAGEN www.qiagenbioinformatics.com). Standard RT-qPCRs were performed using the PowerSYBR-Green PCR Master Mix (Applied Biosystems). The specificity of the amplicons was assessed by visual inspection of the amplification and melting curves of the RT-qPCR and by gel electrophoresis of the PCR products (after 40 thermal cycles with size inspection on 3% agarose). PCR amplification efficiencies of the primer pairs for the 91 target genes and two endogenous control genes were validated as suggested in step 14 and 15 of the protocol of Schmittgen and Livak (2008).

### Expression analysis using high throughput quantitative real-time PCR

Expression analysis used the high throughput system BioMark™ HD (Fluidigm Corporation, Munich, Germany) with dynamic array chips (96.96 GE IFC; Fluidigm) according to the manufacturer’s instruction. Luminescence data recording and processing applied the BioMark Real-Time PCR Analysis Software 3.0.2 (Fluidigm). The overall quality score of the experiment was 0.945. Variation between the chips was low (0.92 to 0.97). C_t_ values of several 96.96 IFC chips were joined with their meta-data in an expression set using the R-package “HT-q-PCR” (Dvinge and Bertone 2009). All C_t_ values below 5 and C_t_ values of genes showing little variation between the samples (with an inter quartile range below 0.6) were discarded.

The relative amount of mRNA molecules was calculated based on the C_t_ value (cycle number at threshold). The cycle threshold was determined with the automatic linear base line setting.

#### Normalization

The genes *VIT_17s0000g10430* encoding glyceraldehyde-3-phosphate dehydrogenase (GAPDH) and *VIT_08s0040g00040* encoding ubiquitin-conjugating enzyme E2 (UBIc; Online resource 3) served as references. These genes had already been successfully applied in other RT-qPCR studies e.g. (Monteiro et al. 2013; Reid et al. 2006; Selim et al. 2012; Upadhyay et al. 2015). Their expression proved to be rank invariant in rachis tissue over clones, locations and growing seasons (as revealed with the function “normalizectdata” of the package “HT-qPCR”). To obtain the ΔC_t_ value, the C_t_ value of each target gene was normalized by subtraction of the mean C_t_ values from the two endogenous reference genes (GAPDH, UBIc). For gene expression comparisons between clones, seasons and vineyard locations, the 2^−ΔΔCt^ value was calculated (Livak and Schmittgen 2001). The relative expression (2^−ΔCt^) of ‘Pinot noir’ clone Gm20-13 at each individual location and season was subtracted from the (2^−ΔCt^) of any other investigated PN clone to standardize.

### Statistics

All statistics employed R-software version 3.5.3 (R Core Team 2013).

Cluster architecture: The environmental impact on each cluster architecture sub-trait was assessed using generalized linear models (GLM) with clone, location, season and the two-way interaction between location and season as explanatory variables. For count data, a GLM with Poisson distribution or (when overdispersed) negative binomial distribution was fitted. For strictly positive continuous responses a Gamma-GLM with log-link or a linear model was applied. Model residuals were visually assessed and dispersion was checked when applicable. Effects were tested using type three analysis of variance and the function “Anova” of the package “car”(Fox and Weisberg 2011), and visualized using the function “alleffects” of the package “effects” (Online resource 4). Estimated marginal means, posthoc tests and pairwise comparisons with compact letter display were calculated for the effect of “clone” on the response while accounting for the effects of “season” and “location” (Table 3) using the functions “emmeans” and “CLD” of the package “emmeans” (Lenth 2019). The significance level was set to 0.05.

Differential gene expression, denoted as fold change (FC), was calculated using the package “limma” (Matthew et al. 2015). First, a design matrix, containing the experimental information for all PN clones at three trial locations and three seasons was generated with the function “model.matrix”. Second, the correlation between technical replicates was estimated with the function “duplicatecorrelation”. The differential gene expression was analyzed by fitting gene-wise linear models using the design matrix, the estimated correlation and the function “lmFit”. To interpret different gene expression, the empirical Bayes method was used to modify the standard errors towards a common value using the “eBayes” function.

Contrast: the log_(2)_ FC (-ΔΔC_t_) for each gene was calculated by the expression difference to the standard clone Gm20-13 (as defined in the contrast matrix) using the function “contrasts.fit” (Online resource 5). The results of relative gene expression were displayed in heatmaps as log_2_ FC (-ΔΔC_t_) with the package “pheatmap” (Kolde 2015). Row-scaled data (gene-wise) and Euclidian distance were used for hierarchical clustering in heatmaps. Expression data of tested genes (log_2_ FC), displayed in box-whisker-plots, were obtained in the same way as stated above but with the model matrix containing additionally the biological replication (Online resource 6).

Variance partition: To estimate the variation in this multilevel gene expression experiment the package “variancePartition” was used with the log_2_ of ΔC_t_. A linear mixed model with the random effects season, location, batch, biological replicate, cluster type, clone and gene pool identified the typical drivers of variance. These were environmental (“season” and “location”), technical (two repeated “batches”), biological (three independent “replicates”), phenotypic (“cluster type”) and genetic (“clone” and “gene pool”, i.e. selection background of ENTAV, Frank, Fr (Freiburg), Gm (Geisenheim) and We (Weinsberg) clones) (Hoffman and Schadt 2016). Correlation between relative test gene expression, expressed as log_(2)_ FC (−ΔC_t_), and cluster architecture sub-trait records of ‘Pinot Noir’ clones for 2015 and 2016 were calculated with Spearman rank correlation using the function “rcorr” from the package “Hmisc”(Harrell Jr 2015) (Table 4, Table 5, Online resources 7, 8).

**Table 4.**
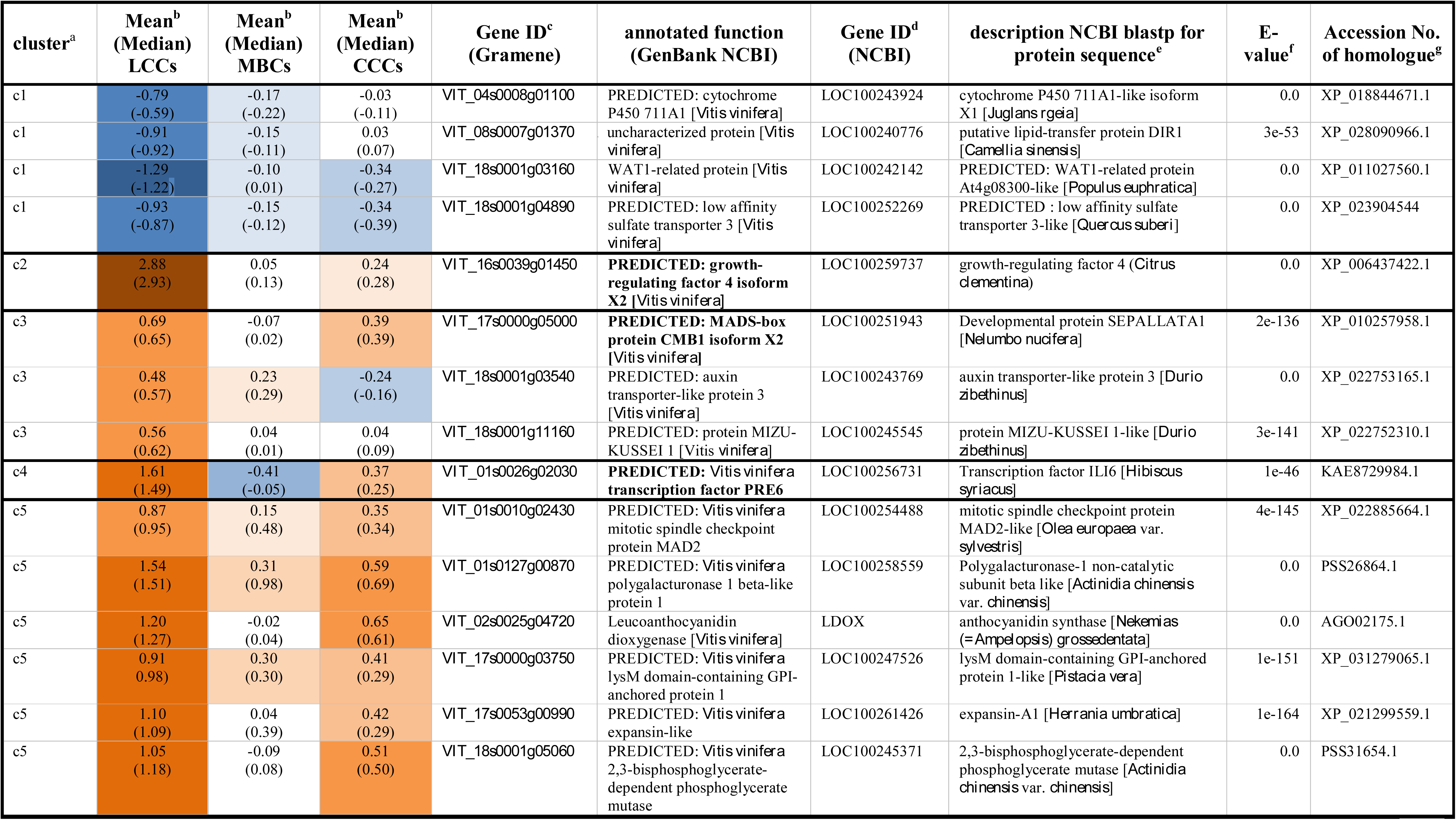
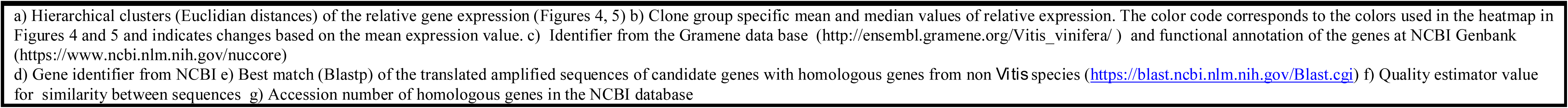
Average gene expression fold change log_(2)_ FC (-ΔΔC_t_) at early fruit development stage (BBCH71) in loosely clustered clones (LCCs), mixed berried clones (MBCs) and compactly clustered clones (CCCs) as compared to the standard ‘Pinot Noir’ clone Gm20-13

**Table 5.**
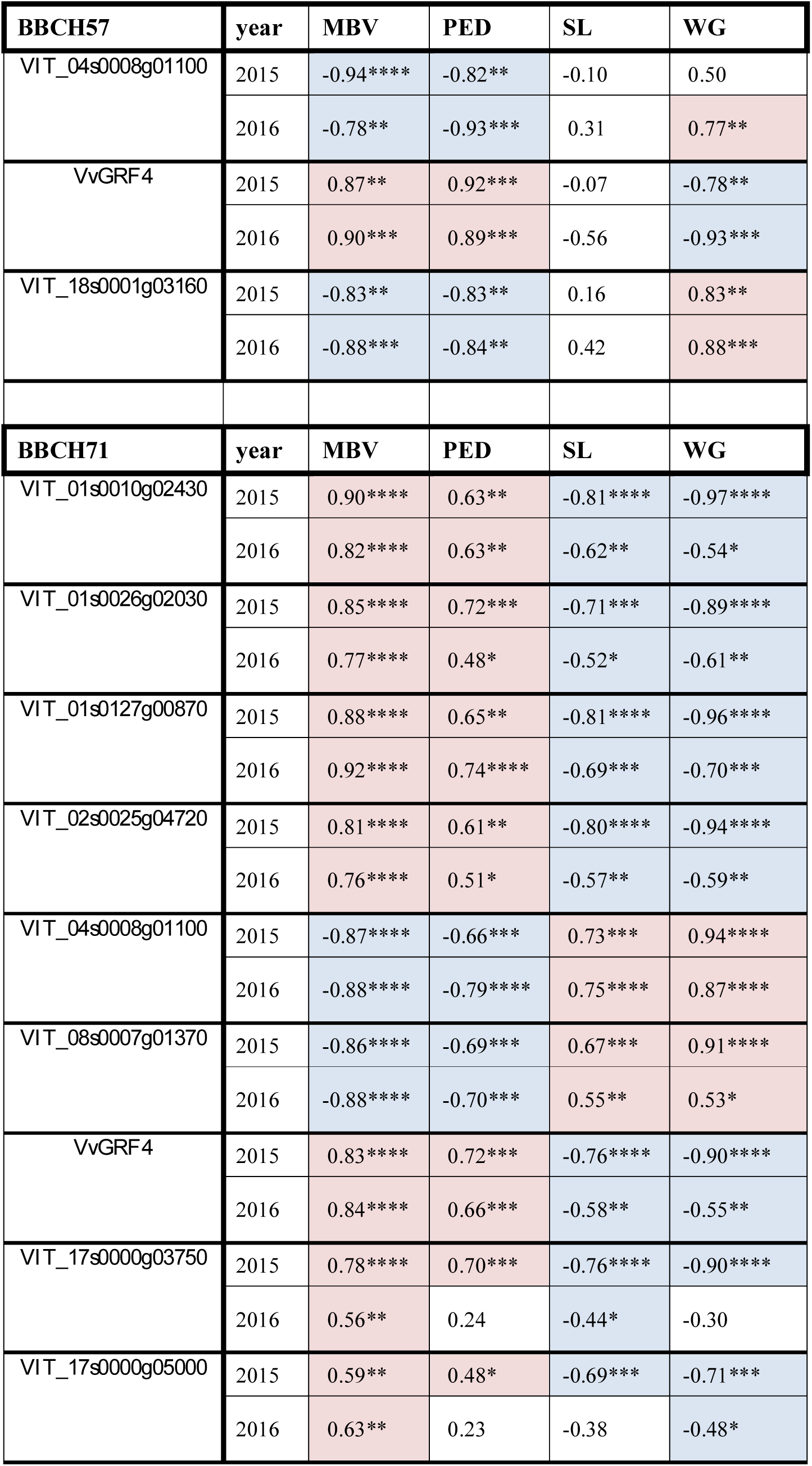
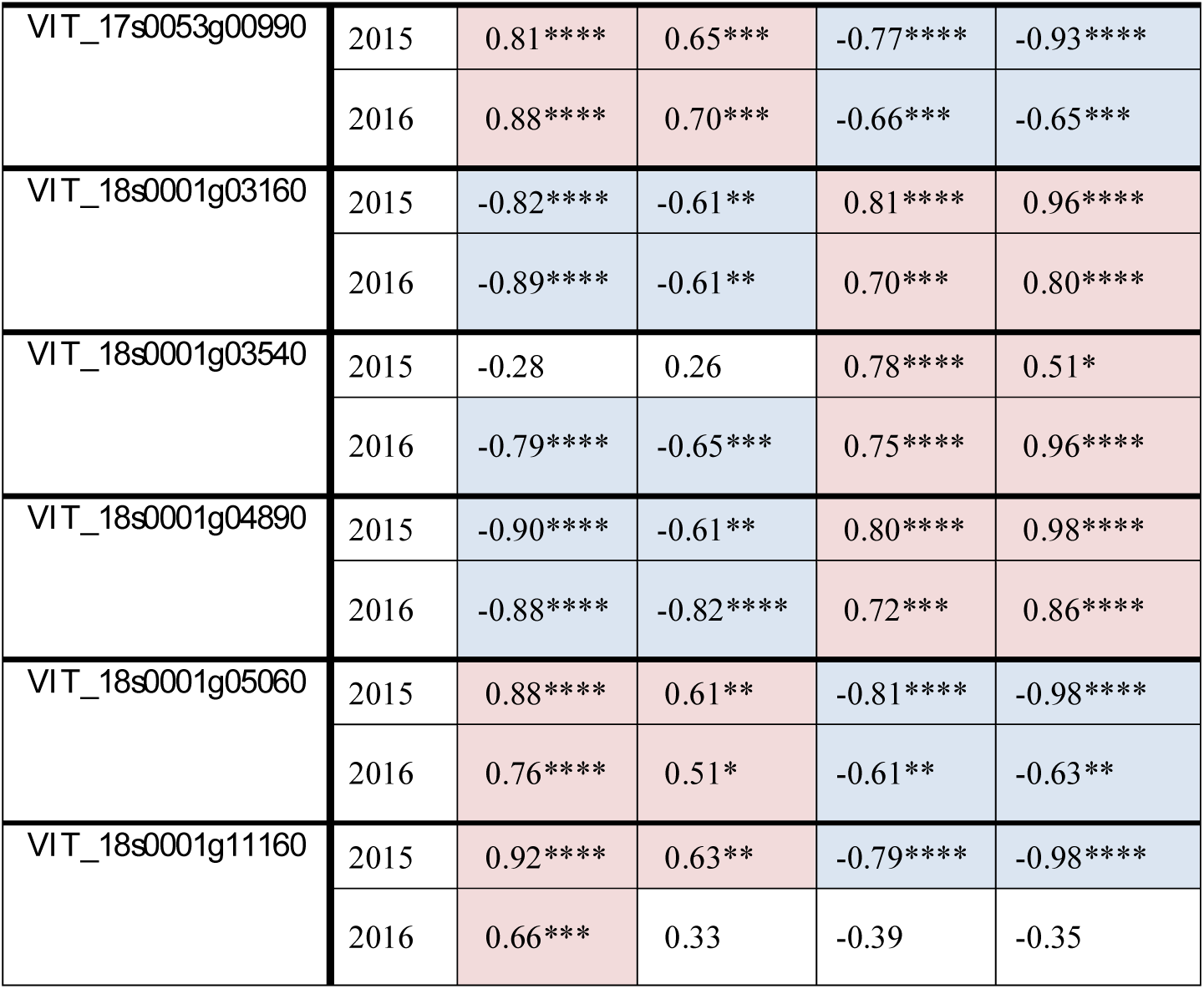
Coefficient of correlation (r) between the relative expression changes of selected genes and key sub-traits of cluster architecture and wood gain (for abbreviations see Table 2). The gene expression relative to (GAPDH) and UBIc (log_(2)_FC) was measured just before flowering (BBCH57) and just after flowering (BBCH71). The results for cluster architecture sub-traits of ‘Pinot Noir’ clones were recorded at ripe grape clusters stage BBCH89. Wood gain was recorded after leaves had fallen (BBCH97). Spearman correlation (r) is significant with *p <0.05, **p <0.01, ***p <0.001 and ****p <0.0001 Positive correlation is labeled in light red, negative correlation in light blue.

## Results

### Cluster architecture features of ‘Pinot Noir’ clones

The typical differences in cluster architecture (CA) exhibited by the investigated (PN) clones at stage BBCH89 (berries ripe for harvest) are depicted in Figure 1.

**Figure 1.**
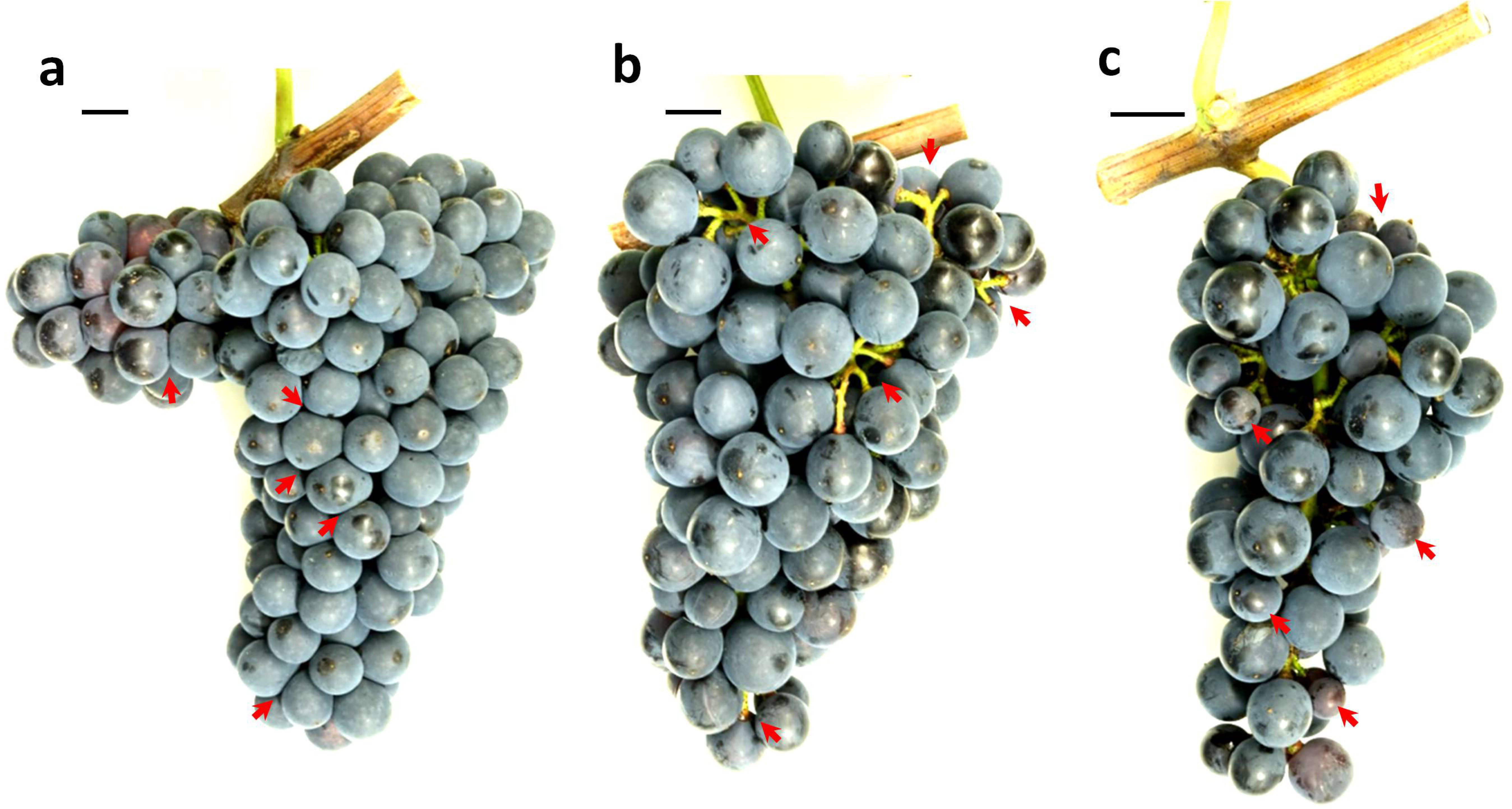
Clones of *Vitis vinifera* cv. ‘Pinot Noir’ with different cluster architecture Phenological stage BBCH89 (berries ripe for harvest) was used for cluster architecture assessment. (a) Compactly clustered clones with non-circular shaped berries due to high pressure between the berries. (b) Loosely clustered clones with visibly extended rachis and pedicel lengths. (c) Clones bearing partially smaller berries leading to reduced compactness (Mixed berried clones). Red arrows highlight the emphasized cluster architecture feature. The size standard depicts 1cm. Developmental stages according to (Lorenz et al. 1995)

The morphological characteristics of ripe bunches were evaluated in 12 PN clones spread over three geographic locations in 2015 and 2016 at BBCH89 (Table 1, Online Resource 1). The PN clones Gm20-13 (MBC) and Frank Charisma (CCC) were represented at all three locations. They allowed to estimate the effect of location and season on cluster architecture traits. The ratio “berry number/rachis length”, and indices CI-12 and CI-18 (Tello and Ibáñez 2014) were applied to categorize the PN clones. Their general visual classification in loose and compact clones (Ruehl et al. 2004) was confirmed and the clones were characterized as three CCC, two MBC and six LCC (Table 1, Table 2). The clone Gm18 remained unclassified due to unstable expression of the sub-traits represented in the indices.

In total, 12 sub-traits of cluster architecture (CA) were evaluated. Ten out of the 12 sub-traits differed significantly between the clones. The lengths of the first rachis internode (I1L) and second rachis internode (I2L) did not vary (Table 3). PN clone Gm20-13 continuously showed low values for sub-traits of CA (small berries, short rachises, i.e. MBV and RL, Table 3). The factors “season” and “location” were evaluated in the clones Gm20-13 and FkCH that were represented at all three locations (Hesse, Baden, Palatinate). “Season” affected the sub-traits berry number (BN), mean berry volume (MBV), total berry volume (TBV), rachis length (RL), shoulder length (SL) and rachis weight (RW). The factor “location” affected the sub-traits cluster weight (CW), mean berry volume (MBV), total berry volume (TBV), rachis length (RL), shoulder length (SL) and rachis weight (RW) (Online resource 4). The values for peduncle lengths (PL), internode sections (L1I, L2I) and pedicel lengths (PED) in Gm20-13 and FkCH were stable and did not differ between locations and seasons (Online resource 4).

To capture the effects of varying vineyard conditions that could affect cluster architecture, the annual wood gain was recorded as indicator of plant vigor (Table 3). The values of clones Gm20-13 and FkCH attained during the seasons 2015 and 2016 differed significantly between the three locations (Online resource 1). The highest wood gain per vine was achieved in Baden (average 1136 g, integrated management), followed by Hesse (average 758 g, integrated management) and Palatinate (average 456 g, vineyard under organic management). Wood gain (WG) was not significantly affected by season (Online resource 4).

Table 3 summarizes the results of the morphometric characterization of the bunches. The loosely clustered clones from Freiburg (Fr12L, Fr13L) and from Weinsberg (WeM1, WeM171, WeM242) shared long rachis lengths and enhanced berry volume. The clones Fr12L, Fr13L and WeM242 showed extended pedicel lengths, as did the loosely clustered clone Gm1-86 from Geisenheim. However, the latter clone (Gm1-86) formed shorter rachises. The compact PN clones in general produced small berries with short pedicels at reduced rachis lengths. The analysis also included mixed berried clones that differed concerning berry volume and berry number in comparison to their co-members from the same clonal selection lines. The PN clones Gm20-13 and Frank Charisma were available in all vineyards and measured over all seasons. This data allowed to investigate the environmental effects (Factor “location” and “season”) on the morphology of bunches as shown in Fig. 2. All the morphometric measurements served to study differential gene expression in association with cluster architecture characteristics.

**Figure 2.**
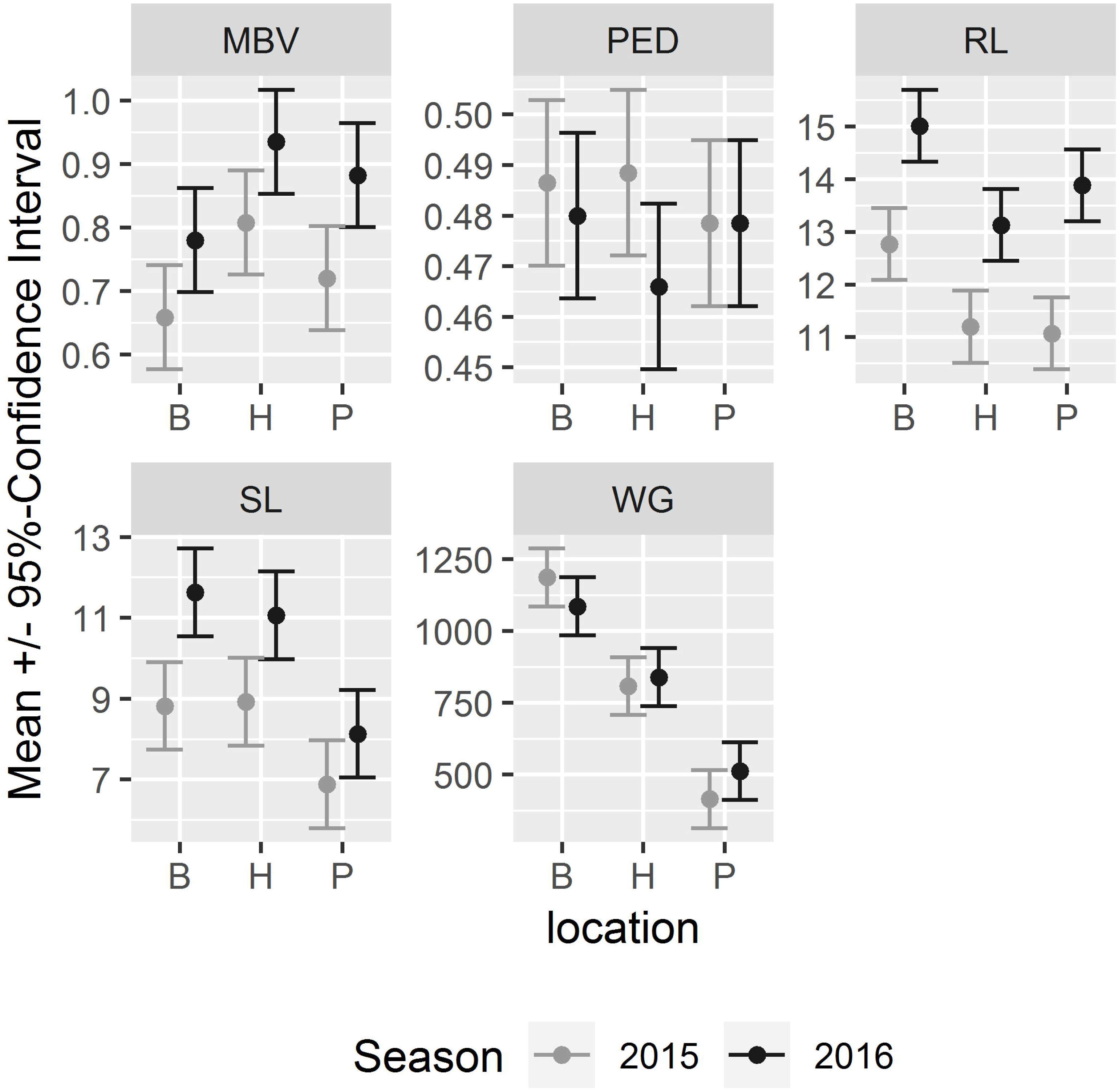
Effects of sampling locations and growing seasons on selected cluster architecture sub-traits and wood gain for the ‘Pinot Noir’ clones Gm20-13 and FkCH. These two clones could be sampled across all seasons and locations. Mean and 95% confidence intervals were estimated with generalized linear models (n = 120). The CA sub-traits rachis length (RL), shoulder length (SL) and mean berry volume (MBV) were clearly influenced by “season”. In contrast, pedicel length (PED) was affected neither by “season” nor by “location” (Online resource 4).

### Identification of regulated genes and expression of *VvGRF4*

Candidate genes were selected from a previous RNA-Seq study and literature references (Online Resource 3). The gene *VvGRF4* was included to check its general implication in cluster compactness in an extended set of PN clones from various selection backgrounds and over different environments. The clone Gm20-13 had a distinct phenotype (small berries, short rachises) and was used as reference to standardize the gene expression data, contrasting its expression with data of those clones that show long rachis features and high berry volume.

Accelerated inflorescence growth of loosely as compared to compactly clustered PN clones just before flowering (BBCH57) and at early fruit set (BBCH71) has been reported (Richter et al. 2017). Hence, these time points were chosen for the expression analysis of 91 genes in the 11 PN clones categorized for their cluster architecture (Figure 3, Table 2). Quantitative Real Time PCR was performed on developed inflorescences (BBCH51) and on young clusters at fruit set (BBCH71).

**Figure 3.**
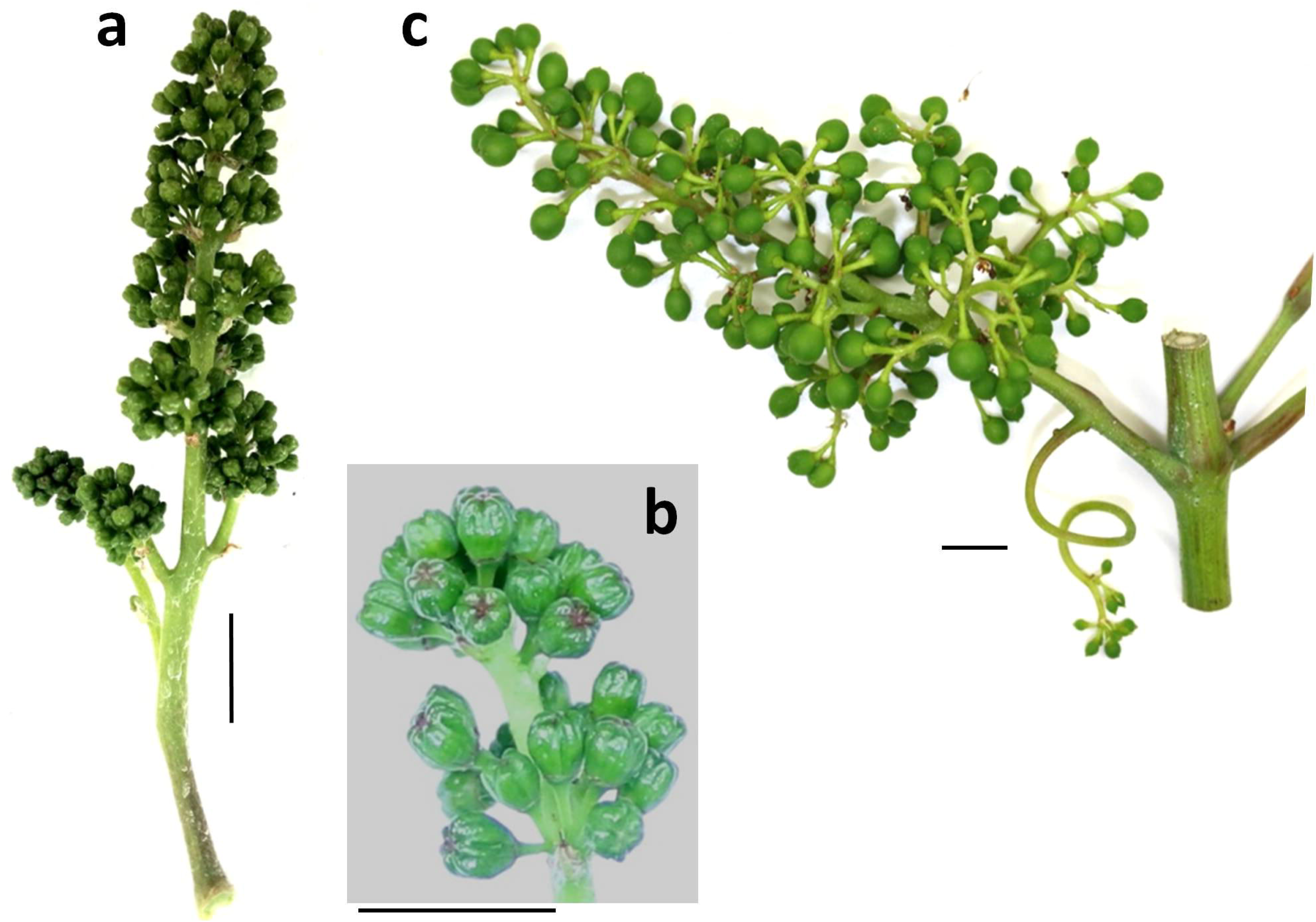
For differential gene expression studies, BBCH57 (a) (just before flowering with still closed flower caps (b)) and BBCH71 (c) (berry set) samples were used for gene expression analysis. For each time point, three biological replicates were collected from different vines. The sampled vines were chosen randomly within a plantation of several hundred individuals of each clonal variant. Only vines without any indication of pathogen infection or physiological disorder were sampled.

In total, 40 genes at BBCH57 and 81 genes at BBCH71 appeared differentially expressed between the PN clones of LCC, MBC or CCC phenotype (Online Resource 5). Out of these, 15 differentially expressed genes were inferred with moderated T-statistics using empirical Bayesian modeling (Smyth 2004). These 15 genes were differentially expressed autonomously, that means independently from “season” and “location”. They included the gene encoding transcription factor *VvGRF4*, as expected from the former study of Rossmann et al (2019), assessed here in a larger clone set. *VvGRF4* was differentially expressed both at BBCH57 and BBCH71. In line with the former results, its activity was high in LCC clones and down-regulated in CCC (Figure 4, Figure 5). The expression of *VvGRF4* in MBCs resembled the pattern seen in CCCs. In addition to *VvGRF4*, two genes (*VIT_04s0008g01100* and *VIT_18s0001g03160*) were consistently differentially active at the early stage of BBCH57 (Fig. 4).

**Figure 4.**
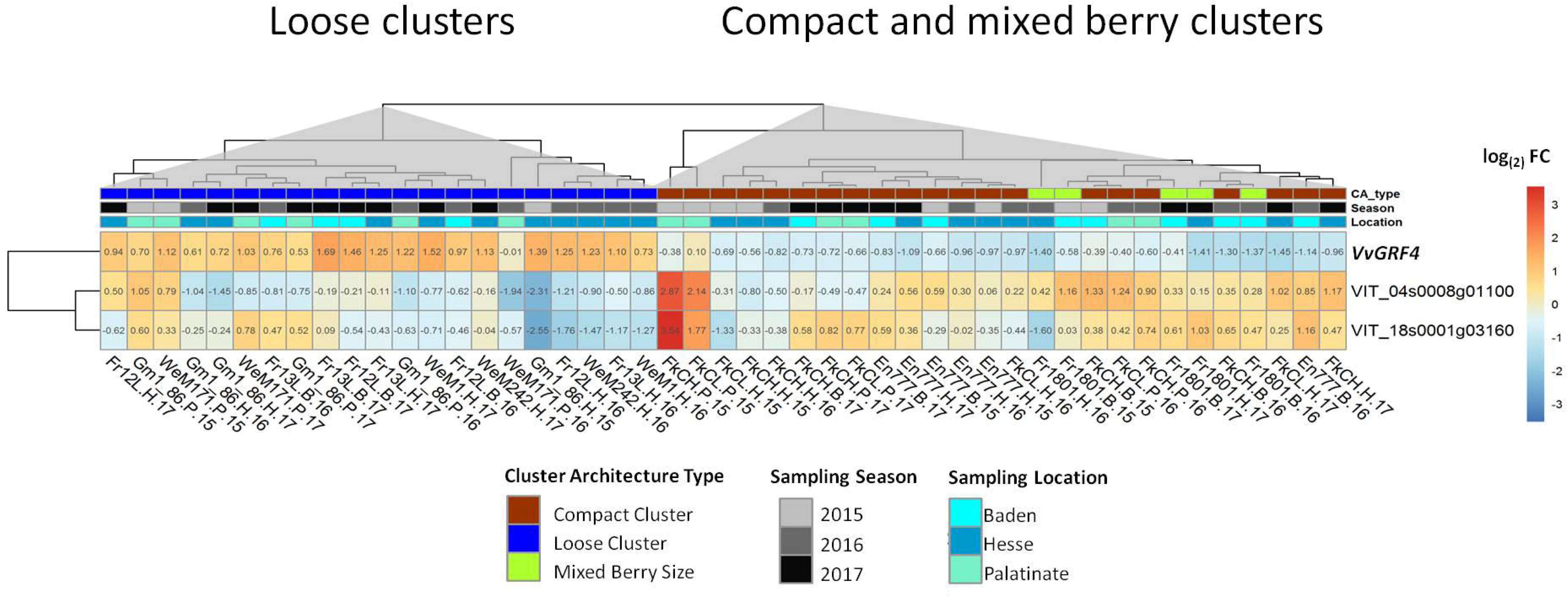
Heatmap of the averaged (three biological and two technical replicates) relative gene expression values as log_(2)_ FC (-ΔΔC_t_) of selected genes at BBCH57. The gene expression relative to the mean of GAPDH and UBIc was analyzed just before flowering (BBCH57) and standardized relative to the PN clone Gm20-13. The rows show the relative expression of the genes. The columns represent the ‘Pinot Noir’ samples. The clones are indicated at the bottom with their abbreviated name, their location (B = Baden, H = Hesse, P = Palatinate) and the year of sampling (15 = 2015, 16 = 2016, 17 = 2017). Hierarchical clustering (based on Euclidian distances) revealed similarities in gene regulation in the PN clones depending on their cluster architecture (CA) type. LCCs are separated from CCCs and MBCs.

**Figure 5.**
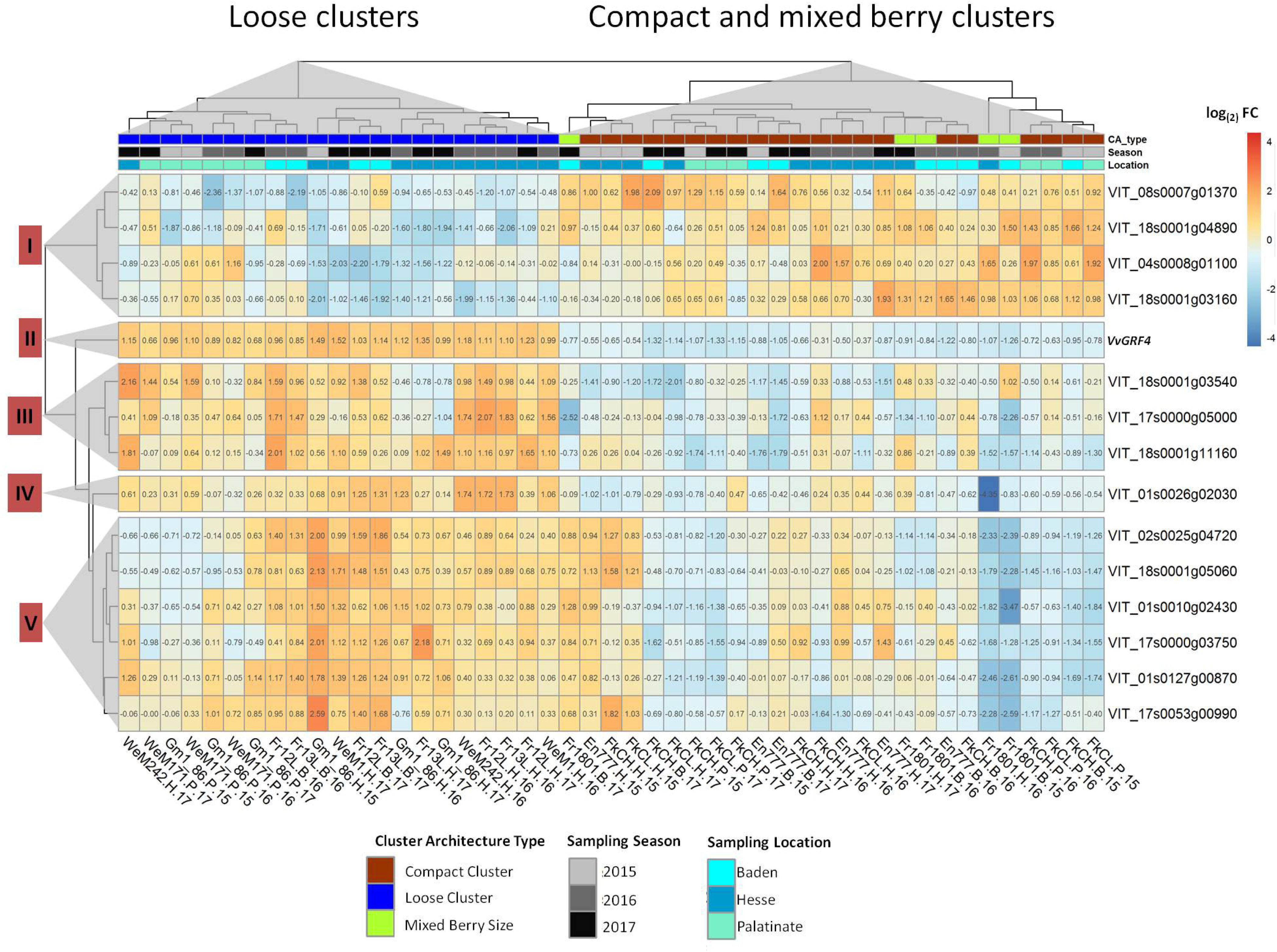
Heatmap of the averaged (three biological and two technical replicates) relative gene expression values as log_(2)_ FC (-ΔΔC_t_) of selected genes at BBCH71. The gene expression relative to the mean of GAPDH and UBIc was analyzed just after flowering (BBCH71) and standardized relative to the PN clone Gm20-13. The rows show the relative expression of the genes. The columns represent the ‘Pinot Noir’ samples. The clones are indicated at the bottom with their abbreviated name, their location (B = Baden, H = Hesse, P = Palatinate) and the year of sampling (15 = 2015, 16 = 2016, 17 = 2017). Hierarchical clustering (based on Euclidian distances) revealed similarities in gene regulation in the PN clones depending on their cluster architecture (CA) type. LCCs are separated from CCCs and MBCs. The genes expression data form five clusters of similar patterns (as indicated by numbers at the left hand side).

After fruit set and begin of fruit development (BBCH71), 11 more genes were found to be differentially expressed between loose and compact PN clones over all seasons and locations. Their regulation reached a higher amplitude as in the young stage (BBCH57). Hierarchical clustering was applied to their expression values. Together with *VvGRF4*, the genes were grouped into five clusters according to their expression patterns (Table 4, Fig.5). The clustering of PN clones showed a clear separation of LCCs from CCCs and MBCs (Fig.5).

In expression cluster I, the transport- and phytohormone related genes *VIT_04s0008g01100, VIT_08s0007g01370, VIT_18s0001g03160* and *VIT_18s0001g04890* were down-regulated in the majority of LCCs, while they showed only little expression changes in most MBCs and CCCs. The gene *VvGRF4* formed a separate cluster II and followed a homogenous differential expression pattern specific to loose resp. compact and mixed-berried clones. It was highly active in LCC clones. Cluster III contained the genes *VIT_17s0000g05000, VIT_18s0001g03540* and *VIT_18s0001g11160*. The products of these genes relate to transcription regulation (transcription factor SEPALLATA1-like), auxin transport and auxin homeostasis. They were up-regulated in most LCCs to a much larger extent, than in CCCs. Cluster IV consists of gene *VIT_01s0026g02030*. It encodes a non-DNA binding basic helix-loop-helix (bHLH) transcription factor PRE6. For this transcription factor gene, the LCCs showed higher expression than the CCCs. The MBCs showed a heterogeneous range of differential expression extending from −4.35 to 0.39. In cluster V, expression patterns showed the highest heterogeneity. The genes *VIT_01s0010g02430, VIT_01s0127g00870, VIT_17s0000g03750* and *VIT_17s0053g00990* encode proteins related to cell wall synthesis or cellular growth. The products of the genes *VIT_02s0025g04720* (LDOX) and *VIT_18s0001g05060 (2, 3-biphosphoglycerate-dependent phosphoglycerate mutase-like)* are associated with pro-anthocyanidin synthesis resp. glycolysis/gluconeogenesis. Few CCC samples showed divergent (up-regulated) gene expression affected by “season” and “location” (e.g. Hesse 2015). The LCC samples from Palatinate showed repression for four genes in cluster V in contrast to the clones from the other locations (Fig. 5). The expression changes are summarized in table 4.

### Variance of gene expression explained by experimental factors

In order to determine to which extent the modulations of gene expression were affected by experimental factors apart from their relationship to cluster architecture, a variance partition analysis was carried out. For all the identified genes, the factor “cluster type” explained a substantial percentage of the variance in gene expression (Fig. 6, Online resource 8). The factors “location” and “season” also showed clear effects.

**Figure 6.**
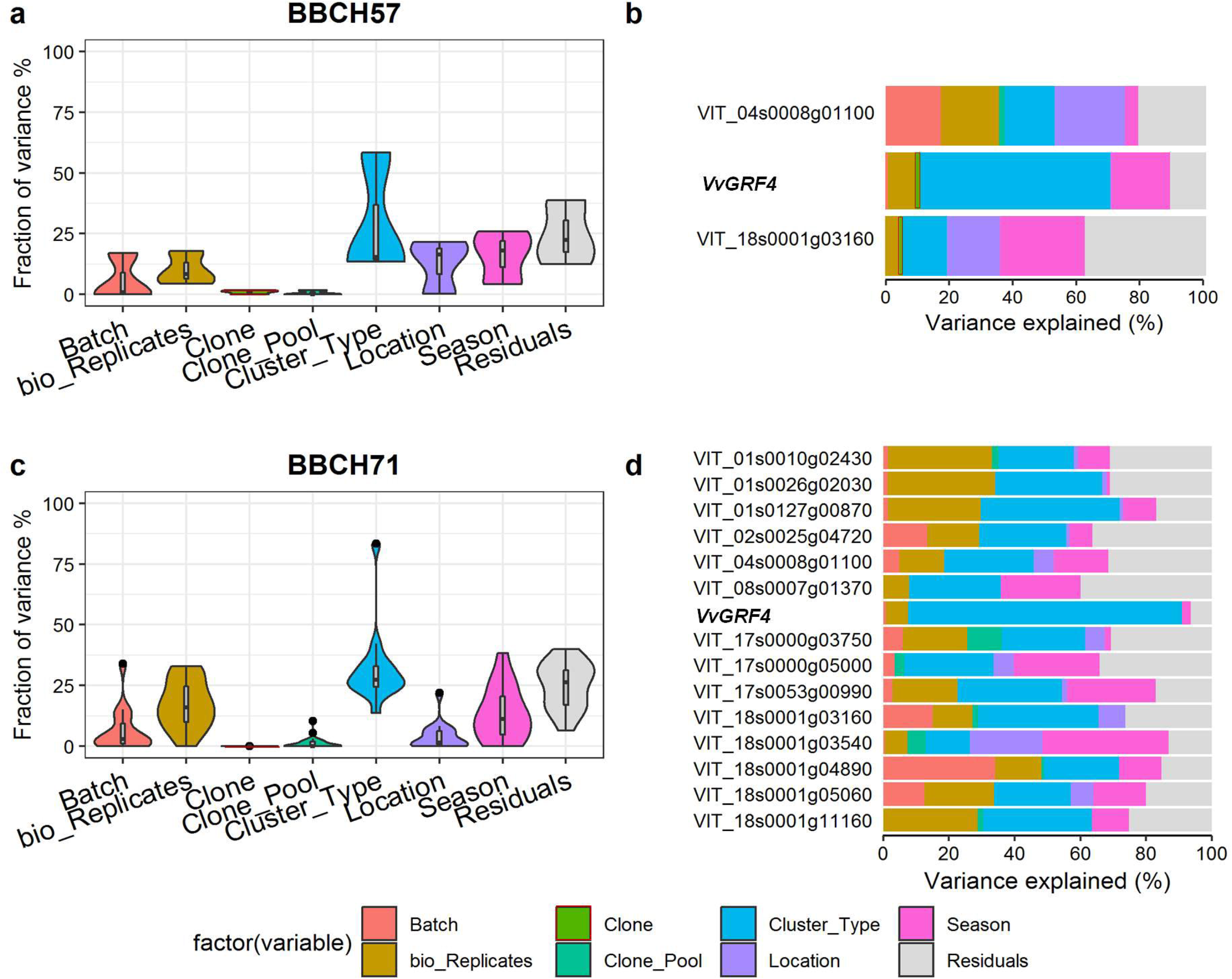
Variance partition analysis using experimental factors to assess the percentage of the explained variance of gene expression. The violin plots (a, c) indicate the explained variances in overall gene expression values log_(2)_ (ΔC_t_) on the y-axis, while the x-axis depicts the factors of variance: cluster type (loose, mixed berried, compact), bio replicates, (biological replicates, n=3), season, batch (technical replicates, n=2), location, gene pool (selection background), clone (11 ‘Pinot Noir’ clones) and the residuals. The bar plots (b, d) depict the amount of variance explained by each factor on the individual gene’s expression.

At the early time point, BBCH57, the main cause of variance for *VvGRF4* was “cluster type” (58% explained variance). For *VIT_18s0001g03160* (a vacuolar auxin transporter) it was “season” (26%). The variance of *VIT_04s0008g01100* (cytochrome P450 711A1) was mainly explained by the factor “location” (22%).

At the later developmental stage, BBCH71, the factor “cluster type” was the major determinant of gene expression variation of almost all 15 investigated genes. The sole exception was *VIT_18s0001g03540* (with only 14% of variance explained by “cluster type”). The variance of *VvGRF4* gene expression was explained to more than 80% by “cluster type”. However, the factor “season” was an important determinant of gene expression variation explaining more than 20% of variance for the genes *VIT_08s0007g01370 VIT_17s0000g05000, VIT_17s0053g00990* and *VIT_18s0001g03540* (Figure 6, Online resource 8).

The gene *VIT_18s0001g04890* was strongly affected by factor “batch” (technical replicates) and the genes *VIT_01s0010g02430, VIT_01s0026g02030, VIT_01s0127g00870* and *VIT_18s0001g11160* varied over the biological replicates (Online resource 8).

### Correlation of gene expression with sub-traits of cluster architecture

At the early stage of BBCH57, the relative expression of *VvGRF4* (log_(2)_ FC) was strongly correlated with the cluster architecture sub-traits mean berry volume (MBV; r= 0.87/0.90) and pedicel length (PED; r= 0.92/0.89) in both years. In contrast, the transcription of genes *VIT_04s0008g01100* and *VIT_18s0001g03160* correlated inversely with MBV and PED (Table 5). There was no significant correlation to shoulder length (SL).

During 2015 and 2016, at developmental stage BBCH71, all selected genes changed expression correlated with at least one of the CA sub-traits mean berry volume (MBV), pedicel length (PED) and shoulder length (SL) (Table 5). Three main trends appeared in both seasons. I) 11 genes with significant correlation to MBV also correlated with PED. Genes with correlation to SL often co-correlated with plant vigor (measured as wood gain, WG). II) The correlations to MBV/PED in general appeared of inverse nature to the correlations observed to SL/WG (Table 5, Online resource 7). III). None of the 15 genes showed any significant correlation with the CA sub-traits berry number (BN), cluster weight (CW) or rachis length (RL) (Online resource 7).

Interestingly, at BBCH71 the correlation of the genes expression with MBV was generally stronger than to PED. This trend was not observed for the three genes regulated at the early stage of BBCH57, where both correlations were about the same in strength. All genes showed expression regulation correlated with the sub-trait shoulder length (SL) in at least one season.

### Correlation of differential gene expression in between the modulated genes

In general, the correlation among the differentially expressed genes was strong, with the sole exception of *VIT_18s0001g03540* (Online resource 6).

Consistent with the gene expression clusters (Figures 4 and 5), the genes that were positively correlated to MBV and PED also correlated positively to the genes of the expression clusters II to V, but negatively to the genes of cluster I (Online resource 7). Vice versa, the genes that correlated negatively to MBV and PED also correlated negatively to all genes in expression clusters II to V, but positively to the genes in cluster I (Online resource 7).

The three genes *VIT_01s0026g02030, VvGRF4* and *VIT_17s0000g05000* encode putative transcription factors. At BBCH57, the expression of *VvGRF4* correlated negatively with the genes differentially expressed at this developmental stage. This negative correlation continued to the later stage. At BBCH71, the expression of the ten regulated genes was always correlated with the transcriptional activity of the three transcription factor genes in the same sense. The three transcription factor genes correlated positively to each other. The gene *VIT_18s0001g04890* correlated with *VIT_17s0000g05000* only during the season of 2015 (Table 6).

**Table 6.** Coefficient of correlation for relative gene expression (log_(2)_FC) between the three putative transcription factors and differentially regulated genes. Spearman correlation (r) is significant with *p <0.05, **p <0.01, ***p <0.001 and ****p <0.0001 Positive correlation is labeled in magenta, negative correlation in light blue.

| BBCH | Gene Id | season | *VIT_01s0026g02030* | *VvGRF4* | *VIT_17s0000g05000* | Annotation<br>According to NCBI<br>blastX results |
| --- | --- | --- | --- | --- | --- | --- |
| 57 | *VIT_04s0008g01100* | 2015 |  | -0.83** |  | cytochrome P450 711A1-like |
| 57 |  | 2016 |  | -0.90*** |  |  |
| 57 | *VIT_18s0001g03160* | 2015 |  | -0.98**** |  | WAT1-related protein |
| 57 |  | 2016 |  | -0.95**** |  |  |
| 71 | *VIT_01s0026g02030* | 2015 |  | 0.97**** | 0.79**** | transcription factor PRE6 |
| 71 |  | 2016 |  | 0.87**** |  |  |
| 71 | *VvGRF4* | 2015 | 0.97**** |  | 0.85**** | growth-regulating factor 4 |
| 71 |  | 2016 | 0.87**** |  | 0.74**** |  |
| 71 | *VIT_17s0000g05000* | 2015 | 0.79**** | 0.85**** |  | SEPALLATA1-like protein |
| 71 |  | 2016 | 0.89**** | 0.74**** |  |  |
| 71 | *VIT_01s0010g02430* | 2015 | 0.95**** | 0.93**** | 0.70*** | mitotic spindle checkpoint protein<br>MAD2-like |
| 71 |  | 2016 | 0.92**** | 0.97**** | 0.72*** |  |
| 71 | *VIT_01s0127g00870* | 2015 | 0.92**** | 0.95**** | 0.73*** | polygalacturonase 1 beta-like protein |
| 71 |  | 2016 | 0.82**** | 0.96**** | 0.66*** |  |
| 71 | *VIT_02s0025g04720* | 2015 | 0.88**** | 0.92**** | 0.79**** | anthocyanidin synthase |
| 71 |  | 2016 | 0.98**** | 0.92**** | 0.83**** |  |
| 71 | *VIT_17s0000g03750* | 2015 | 0.89**** | 0.94**** | 0.81**** | lysM domain-containing GPI-anchored<br>protein 1-like |
| 71 |  | 2016 | 0.89**** | 0.83**** | 0.84**** |  |
| 71 | *VIT_17s0053g00990* | 2015 | 0.90**** | 0.92**** | 0.75**** | alpha-expansin |
| 71 |  | 2016 | 0.86**** | 0.97**** | 0.68*** |  |
| 71 | *VIT_18s0001g05060* | 2015 | 0.90**** | 0.92**** | 0.71*** | bisphosphoglycerate-dependent<br>phosphoglycerate mutase-like |
| 71 |  | 2016 | 0.97**** | 0.89**** | 0.81**** |  |
| 71 | *VIT_18s0001g11160* | 2015 | 0.92**** | 0.92**** | 0.69*** | protein MIZU-KUSSEL 1-like |
| 71 |  | 2016 | 0.89**** | 0.86**** | 0.89**** |  |
| 71 | *VIT_04s0008g01100* | 2015 | -0.90**** | -0.87**** | -0.60** | cytochrome P450 711A1-like |
| 71 |  | 2016 | -0.67*** | -0.74**** | -0.42* |  |
| 71 | *VIT_08s0007g01370* | 2015 | -0.90**** | -0.86**** | -0.56** | putative lipid-transfer protein DIR1 |
| 71 |  | 2016 | -0.72*** | -0.88**** | -0.64** |  |
| 71 | *VIT_18s0001g03160* | 2015 | -0.89**** | -0.92**** | -0.74**** | WAT1-related protein |
| 71 |  | 2016 | -0.91**** | -0.89**** | -0.74**** |  |
| 71 | *VIT_18s0001g03540* | 2015 | -0.25 | -0.35 | -0.39 | auxin influx carrier (AUX1 LAX<br>family) |
| 71 |  | 2016 | -0.56** | -0.57** | -0.38 |  |
| 71 | *VIT_18s0001g04890* | 2015 | -0.91**** | -0.91**** | -0.68*** | low affinity sulfate transporter 3-like |
| 71 |  | 2016 | -0.62** | -0.72*** | -0.42 |  |

## Discussion

This study aimed to identify genes involved in the determination of loose cluster architecture in ‘Pinot Noir’ (PN) clones and to check the generality of the implication of *VvGRF4*, recently identified as an important regulator of cluster architecture (Rossmann et al., 2019). In this context, 12 different PN clones with varying types of cluster architecture were studied in detail for their sub-traits that determine the overall phenotype. Enlarging the range of cluster architecture types investigated in the previous study (that was conducted only on two loose and two compact PN clones), the cluster type of “mixed berried-clones” was added to compact and loosely clustered PN clones. The developmental stage of beginning berry formation was investigated for gene regulation as a relevant stage for the constitution of final berry size.

The PN clones were studied over two (for phenotyping) resp. three (for gene expression analysis) seasons in three different environments to identify sub-traits of cluster architecture and their responsible genes that operate independently from local and seasonal conditions. As reviewed in Grishkevich and Yanai (2013), the phenotype of an organism is determined by a combination of its genotype (G), environment (E) and their interaction (G×E). Therefore, it is desirable to dispose of high numbers of clonal individuals spread over several locations. However, for perennial crops like grapevine, this requirement is difficult to fulfil. Establishment of controlled vineyards raised from certified plant material with ample material to allow random sampling is time-consuming and causes high costs. The PN clones in this study needed to be grown in homogeneous plots and grafted on the same rootstock cultivar to avoid transcriptomic shifts in the scion and significant influence on yield and vigor by the rootstock (Chitarra et al. 2017). The experimentation here was therefore restricted to clonal material available at the collaborating nurseries and the cultivar repository at the Institute (Geilweilerhof). The three plantations were managed differently (organic viticulture at Geilweilerhof, integrated management at the nurseries), a fact which should delimit the identification of genetic components to those that operate autonomously from environmental conditions.

The clones FkCH and Gm20-13 were present at all three locations and allowed the estimation of the environmental impact on the cluster architecture phenotype and the differential expression of involved genes. WG (wood gain) and RL (rachis length) were strongly affected by the environmental conditions, while especially PED (pedicel length) differentiated these loose (Gm20-13) and compact (FkCH) clones in a stable way, independent from the vineyard location.

In the investigation of cluster architecture characteristics over all clones, the sub-traits MBV (mean berry volume), berry number (BN), RL (rachis length), SL (shoulder length), and PED (pedicel lengths) emerged as most relevant for the expression of overall cluster architecture.

The differential expression of candidate genes was studied exactly at the time when the phenotypic trait changes between loosely clustered clones and compactly clustered clones. It corresponds to a phase of accelerated rachis growth occurring in young, developing loose clusters in comparison to compact and mixed berried clones (Richter et al. 2017). This results in enhanced cell numbers of pedicels (Rossmann et al. 2019)

For gene expression analysis, the standard clone Gm20-13 with its distinct phenotype of characteristically small berries and short rachis was used as reference to standardize gene expression in the 12 PN clones. The work focused on 15 genes that were differentially expressed during cluster development under all different environmental conditions. These included the gene encoding VvGRF4 and confirmed its importance in the regulation of cluster phenotype (Online resource 6). However, the regulation of these genes was affected by environmental and experimental fluctuations. Nevertheless, analysis of the part of variance explained by sampling and environmental factors, in addition to the cluster architecture, confirmed a prominent percentage of their expression variation as linked to the bunch compactness phenotype (Figure 6).

From 91 genes tested, three genes at BBCH57, and 12 more genes at BBCH71 exhibited regulation linked to cluster architecture over all locations and seasons. Samples from the pre-bloom time showed less variation related to cluster architecture and more variation due to location and season (Figure 6, Online resource 8), than at the later stage. This corresponds to the observation of Dal Santo et al. (2018), who identified the factors “developmental stage”, “season” and “location” to affect the overall transcriptional variation over three seasons in two grapevine cultivars.

At the early stage of BBCH57, the expression of *VvGRF4* was already augmented in the loosely clustered clones, and –inversely– repressed in compact and mixed berried clones. A subtle modulation was observed in the genes *VIT_04s0008g01100* and *VIT_18s0001g03160* at this point. These two genes are members of cluster I of the gene regulatory groups of the later stage (BBCH71). Overall, they still showed moderate expression changes at fruit set, with a more explicit up-regulation in compact and mixed berried clones. *VIT_18s0001g03160* is annotated as a WAT1 (“walls are thin”) encoding gene, a vacuolar transporter of auxin characterized in *Arabidopsis* (Ranocha et al. 2013). The gene *VIT_04s0008g01100* encodes a homolog to cytochrome 450 711A1, a monooxygenase involved in the metabolism of strigolactones (conversion of carlactone to carlactonic acid). Its function has been identified in the *MAX1* mutation in Arabidopsis, which shows increased axillary growth. MAX1 suppresses shoot branching in *Arabidopsis* (Abe et al. 2014). This study here indicates additional or different functions of these genes in grapevine. The cluster I genes with down-regulation in loose clusters also encompass *VIT_18s0001g04890*, annotated as a sulfate transporter. The two genes, *VIT_18s0001g04890* and *VIT_18s0001g03160*, have been described to be repressed in ‘Garnacha Tinta’ clones with larger berries (Grimplet et al. 2017). This is in line with our results here for ‘Pinot Noir’ clones, as high MBV (mean berry volume) corresponds to large berries, which are an important determinant for loose clusters.

Apart from the gene encoding VvGRF4, which was definitely higher expressed in the LCC clones at BBCH71 (Online resource 6), the genes with autonomous up-regulation, particularly in LCCs, included *VIT_17s0000g05000*. This gene encodes a SEPALLATA 1-like developmental regulator. It has probable transcription factor function and is known to be part of the network that regulates flower development in *Arabidopsis* where it prevents indeterminate growth of the flower meristem (Pelaz et al. 2000). Recently, Palumbo et al. (2019) reported *VIT_17s0000g05000* as homeotic gene associated to whorl differentiation in grapevine during the period of pre-anthesis on to post-fertilization. In this study, the mixed berried PN clones Gm20-13 (standard with short rachis) and Fr1801 (long rachis) showed the highest differential expression of *VIT_17s0000g05000* in all three seasons and at all available locations at fruit stet, but not at the pre-flowering stage. These two clones vary mainly in rachis lengths. Thus *VIT_17s0000g05000* represents an interesting candidate controlling rachis length manifestation. It is distinctly down-regulated in the long rachis phenotype (Fr1801). Possibly, this SEPALLATA-homolog is not only involved in flower formation, but also later on in frutescence development.

In addition to auxin transport functions (*VIT_18s0001g03540*) and auxin homeostasis (*VIT_18s0001g11160, Mizu-KusselI* (Moriwaki et al. 2011)), expression of the transcription factor gene *PRE6* was significantly enhanced in LCCs. It belongs to the atypical bHLH transcription factors with no direct DNA binding ability that mediate auxin, brassinosteroid and light signaling and affect photomorphogenesis. A homolog from rice called ILI1 (increased lamina inclination 1) increased cell elongation (Zhang et al. 2009). Cell elongation may well contribute to important cluster features such as rachis length and shoulder length. The further genes with up-regulation, particularly in loose clustered PN clones, encompass functions involved in cell wall extension (*VIT_17s0053g00990*), cell size (*VIT_01s0127g00870)* and cell division *(VIT_01s0010g02430)*. The gene *VIT_17s0053g00990* encodes α-expansin, that was found up-regulated in rapidly growing grape berries and enlarges cell size (Suzuki et al. 2015).

Interestingly, the two clones Gm1-86 and Gm20-13, both originating from the selection line at Geisenheim, differ by berry number (Table 3). This phenotypic difference corresponds to an elevated expression of *VvGRF4* and the gene *VIT_01s0127g00870 (Vitis vinifera* polygalacturonase 1 beta-like protein 1) in Gm1-86, the clone with higher berry numbers. The activity of *VIT_18s0001g04890* (a sulfate transporter) was reduced in Gm1-86 as compared to Gm20-13. However, this gene, encoding a sulfate transporter, showed high variability (34%) within the technical replicates (Online resource 8).

In a previous genetic study, QTL clusters associated with loose bunch architecture were localized in an completely independent genetic background (Richter et al., 2018). Arrays of overlapping QTL regions were found on seven chromosomes, including chromosome 1 and 17. Interestingly, the three genes *VIT_01s0026g02030 (PRE6), VIT_17s0000g05000* (*SEPI*), *VIT_17s0053g00990* (encoding α-expansin), associated to the different cluster architecture characteristics found here for PN clones, are located in QTL areas. Two of them are transcription factors that may have a comprehensive function, which needs to be further investigated.

## Conclusions

This study investigated gene expression in ‘Pinot Noir’ clones of different cluster architecture grown at several locations over three seasons. It revealed 15 genes that were differentially regulated between loosely and compactly clustered clones, independent from year and location (or any other environmental variation encountered). It confirmed the important role of *VvGRF4* in the regulation of cluster architecture in ‘Pinot Noir’. It newly identified two more transcription factor genes, a *SEPALLATA1* homolog and a homolog of *PRE6*, that are more active in the loosely clustered than in the compact bunch type clones. Compared to recent literature, these regulator genes may have new or additional functions in affecting the structure of the ‘Pinot Noir’ grapevine bunch. Furthermore, genes involved in auxin metabolism, cellular growth and transport were found to be regulated. A gene homolog of CYP711A1, encoding an enzyme of strigolactone metabolism, was also involved. Strigolactones function as shoot branching inhibitors (Gomez-Roldan et al. 2008). This gene is repressed in loose clusters, possibly releasing some inhibition, and thus seems to contribute to the loose-clustered phenotype in grapes.

These results improve the basic knowledge on grapevine cluster phenotype. This study revealed several major regulators of cluster architecture in ‘Pinot Noir’, which deserve further attention and functional studies. Studies on the genetic diversity of such major regulator genes in other *V. vinifera* cultivars will show if they are applicable as molecular tools for breeding of advantageous loosely clustered grapevine cultivars with improved resilienceto*Botrytiscinerea*.

## Supporting information

Online resource 1

Online resource 2

Online resource 3

Online resource 4

Online resource 5

Online resource 6

Online resource 7

Online resource 8

## Declarations

## Acknowledgements

This work was funded by the “Federal Program for Ecologic Landuse and other forms of Sustainable Agriculture” (Bundesprogramm Ökologischer Landbau und andere Formen nachhaltiger Landwirtschaft, BÖLN) of BLE (Bundesanstalt für Landwirtschaft und Ernährung) Federal Office for Agriculture and Food under the title “MATA-Molekulare Analyse der Traubenarchitektur” (Molecular analysis of cluster architecture) FKZ 2811NA056 (www.ble.de). We thank Daniel Zendler for fruitful discussion and Margareta Schneider for technical help. We wish to thank the grapevine nurseries Reben Sibbus GmbH, Sasbach-Jechtingen Germany and Antes Viticulture & Grafting GbR, Heppenheim Germany for providing access to their clonal material and maintaining the trial fields in Baden and Hesse.

## Contributions

EZ and RR designed the study. EZ acquired funding and supervised the work. RR performed the experiments, measurements and calculations. SR and KT contributed RNA sequencing data.

DG provided statistical expertise. RT provided plant material, infrastructure and special advice. RR and EZ wrote the paper. All authors read the manuscript.

Supplementary information accompanies the manuscript on the Theoretical and applied Genetics website.

## References

Abe S, Sado A, Tanaka K, Kisugi T, Asami K, Ota S, Kim HI, Yoneyama K, Xie X, Ohnishi T, Seto Y, Yamaguchi S, Akiyama K, Yoneyama K, Nomura T (2014) Carlactone is converted to carlactonoic acid by MAX1 in Arabidopsis and its methyl ester can directly interact with AtD14 in vitro. Proc Natl Acad Sci U S A 111:18084–18089

Alonso-Villaverde V, Boso S, Luis Santiago J, Gago P, Martínez M-C (2008) Relationship Between Susceptibility to Botrytis Bunch Rot and Grape Cluster Morphology in the *Vitis vinifera* L. Cultivar Albariño. International Journal of Fruit Science 8:251–265

Ban Y, Mitani N, Sato A, Kono A, Hayashi T (2016) Genetic dissection of quantitative trait loci for berry traits in interspecific hybrid grape (*Vitis labruscana* × *Vitis vinifera*). Euphytica 211:295–310

Becker T, Knoche M (2012) Water induces microcracks in the grape berry cuticle. Vitis 51:141–142

Bleyer K (2001) Klonzüchtung beim Blauen Spätburgunder. Rebe & Wein 11:22–26

Chitarra W, Perrone I, Avanzato CG, Minio A, Boccacci P, Santini D, Gilardi G, Siciliano I, Gullino ML, Delledonne M, Mannini F, Gambino G (2017) Grapevine Grafting: Scion Transcript Profiling and Defense-Related Metabolites Induced by Rootstocks. Frontiers in Plant Science 8, 654

Citri A, Pang ZPP, Sudhof TC, Wernig M, Malenka RC (2012) Comprehensive qPCR profiling of gene expression in single neuronal cells. Nature Protocols 7:118–127

Correa J, Mamani M, Munoz-Espinoza C, Laborie D, Munoz C, Pinto M, Hinrichsen P (2014) Heritability and identification of QTLs and underlying candidate genes associated with the architecture of the grapevine cluster (*Vitis vinifera* L.). Theoretical and Applied Genetics 127:1143–1162

Dal Santo S, Zenoni S, Sandri M, De Lorenzis G, Magris G, De Paoli E, Di Gaspero G, Del Fabbro C, Morgante M, Brancadoro L, Grossi D, Fasoli M, Zuccolotto P, Tornielli GB, Pezzotti M (2018) Grapevine field experiments reveal the contribution of genotype, the influence of environment and the effect of their interaction (G×E) on the berry transcriptome. The Plant Journal 93:1143–1159

De Lorenzis G, Squadrito M, Rossoni M, Di Lorenzo GS, Brancadoro L, Scienza A (2017) Study of intra-varietal diversity in biotypes of Aglianico and Muscat of Alexandria (*Vitis vinifera* L.) cultivars. Australian Journal of Grape and Wine Research 23:132–142

Di Genova A, Almeida AM, Munoz-Espinoza C, Vizoso P, Travisany D, Moraga C, Pinto M, Hinrichsen P, Orellana A, Maass A (2014) Whole genome comparison between table and wine grapes reveals a comprehensive catalog of structural variants. BMC Plant Biol 14:7 doi: 10.1186/1471-2229-14-7

Döring J, Frisch M, Tittmann S, Stoll M, Kauer R (2015) Growth, Yield and Fruit Quality of Grapevines under Organic and Biodynamic Management. PLOS ONE 10:e0138445

Dry PR, Longbottom ML, McLoughlin S, Johnson TE, Collins C (2010) Classification of reproductive performance of ten winegrape varieties. Australian Journal of Grape and Wine Research 16:47–55

Dvinge H, Bertone P (2009) HTqPCR: high-throughput analysis and visualization of quantitative real-time PCR data in R. Bioinformatics 25:3325–3326

Fanizza G, Lamaj F, Costantini L, Chaabane R, Grando MS (2005) QTL analysis for fruit yield components in table grapes (Vitis vinifera). Theoretical and Applied Genetics 111:658–664

FAOSTAT (2016) http://www.fao.org/faostat/en/#data/. Value of Agricultural Production. Food and Agriculture Organization of the United Nations, last accessed Feb 2, 2020.

Forneck A, Benjak A, Ruehl E (2009) Grapevine (*Vitis* ssp.): Example of Clonal Reproduction in Agricultural Important Plants. pp 581–598

Fox J, Weisberg S (2011) An {R} Companion to Applied Regression, Second edn. [SAGE], Thousand Oaks CA

Gabler FM, Smilanick JL, Mansour M, Ramming DW, Mackey BE (2003) Correlations of morphological, anatomical, and chemical features of grape berries with resistance to *Botrytis cinerea*. Phytopathology 93:1263–1273

Gomez-Roldan V, Fermas S, Brewer PB, Puech-Pages V, Dun EA, Pillot JP, Letisse F, Matusova R, Danoun S, Portais JC, Bouwmeester H, Becard G, Beveridge CA, Rameau C, Rochange SF (2008) Strigolactone inhibition of shoot branching. Nature 455:189–U122

Grimplet J, Ibáñez S, Baroja E, Tello J, Ibáñez J (2019) Phenotypic, Hormonal, and Genomic Variation Among *Vitis vinifera* Clones With Different Cluster Compactness and Reproductive Performance. Frontiers in plant science 9, 1917

Grimplet J, Tello J, Laguna N, Ibáñez J (2017) Differences in Flower Transcriptome between Grapevine Clones Are Related to Their Cluster Compactness, Fruitfulness, and Berry Size. Frontiers in Plant Science 8:17

Grishkevich V, Yanai I (2013) The genomic determinants of genotype x environment interactions in gene expression. Trends in Genetics 29:479–487

Harrell Jr FE (2015) Package ‘Hmisc’. CRAN2018:https://cran.r-project.org/web/packages/Hmisc/Hmisc.pdf.

Hed B, Ngugi HK, Travis JW (2009) Relationship Between Cluster Compactness and Bunch Rot in Vignoles Grapes. Plant Disease 93:1195–1201

Hed B, Ngugi HK, Travis JW (2010) Use of Gibberellic Acid for Management of Bunch Rot on Chardonnay and Vignoles Grape. Plant Disease 95:269–278

Herzog K, Wind R, Töpfer R (2015) Impedance of the Grape Berry Cuticle as a Novel Phenotypic Trait to Estimate Resistance to *Botrytis cinerea*. Sensors 15:12498–12512

Hoffman GE, Schadt EE (2016) variance Partition: interpreting drivers of variation in complex gene expression studies. BMC Bioinformatics 17:483

Houel C, Chatbanyong R, Doligez A, Rienth M, Foria S, Luchaire N, Roux C, Adiveze A, Lopez G, Farnos M, Pellegrino A, This P, Romieu C, Torregrosa L (2015) Identification of stable QTLs for vegetative and reproductive traits in the microvine (*Vitis vinifera* L.) using the 18 K Infinium chip. BMC Plant Biology 15

Keulemans W, Bylemans D, De Coninck B (2019) Farming without plant protection products doi: 10.2861/05433 PE 634.416 ISBN: 978-92-846-3993-9

Kolde R (2015) pheatmap: Pretty Heatmaps. R package version 1.0. 8

Konrad H, Lindner B, Bleser E, Rühl EH (2003) Strategies in the genetic selection of clones and the preservation of genetic diversity within varieties. 603 edn. International Society for Horticultural Science (ISHS), Leuven, Belgium, pp 105–110

Lenth R (2019) emmeans: Estimated Marginal Means, aka Least-Squares Means (Version 1.3.4)

Li M, Klein LL, Duncan KE, Jiang N, Chitwood DH, Londo JP, Miller AJ, Topp CN (2019) Characterizing 3D inflorescence architecture in grapevine using X-ray imaging and advanced morphometrics: implications for understanding cluster density. Journal of Experimental Botany 70 (21): 6261–6276 https://doi.org/10.1093/jxb/erz394

Livak KJ, Schmittgen TD (2001) Analysis of relative gene expression data using real-time quantitative PCR and the 2(T)(-Delta Delta C) method. Methods 25:402–408

Lorenz DH, Eichhorn KW, Bleiholder H, Klose R, Meier U, Weber E (1995) Growth Stages of the Grapevine: Phenological growth stages of the grapevine (*Vitis vinifera* L. ssp. *vinifera*)—Codes and descriptions according to the extended BBCH scale. Australian Journal of Grape and Wine Research 1:100–103

Marguerit E, Boury C, Manicki A, Donnart M, Butterlin G, Nemorin A, Wiedemann-Merdinoglu S, Merdinoglu D, Ollat N, Decroocq S (2009) Genetic dissection of sex determinism, inflorescence morphology and downy mildew resistance in grapevine. Theoretical and Applied Genetics 118:1261–1278

Matthew ER, Belinda P, Di W, Yifang H, Charity WL, Wei S, Gordon KS (2015) limma, powers differential expression analyses for RNA-sequencing and microarray studies. Nucleic Acids Research 43:47

Maul E (2019) Vitis International Variety Catalogue. www.vivc.de

Mejia N, Gebauer M, Munoz L, Hewstone N, Munoz C, Hinrichsen P (2007) Identification of QTLs for seedlessness, berry size, and ripening date in a seedless x seedless table grape progeny. American Journal of Enology and Viticulture 58:499–507

Migicovsky Z, Sawler J, Gardner KM, Aradhya MK, Prins BH, Schwaninger HR, Bustamante CD, Buckler ES, Zhong G-Y, Brown PJ, Myles S (2017) Patterns of genomic and phenomic diversity in wine and table grapes. Horticulture Research 4:17035

Molitor D, Behr M, Hoffmann L, Evers D (2012) Impact of Grape Cluster Division on Cluster Morphology and Bunch Rot Epidemic. American Journal of Enology and Viticulture 63:508

Molitor D, Junk J, Evers D, Hoffmann L, Beyer M (2014) A High-Resolution Cumulative Degree Day-Based Model to Simulate Phenological Development of Grapevine. American Journal of Enology and Viticulture 65:72–80

Monteiro F, Sebastiana M, Pais MS, Figueiredo A (2013) Reference gene selection and validation for the early responses to downy mildew infection in susceptible and resistant *Vitis vinifera* cultivars. PLoS One 8:e72998

Moriwaki T, Miyazawa Y, Kobayashi A, Uchida M, Watanabe C, Fujii N, Takahashi H (2011) Hormonal regulation of lateral root development in Arabidopsis modulated by MIZ1 and requirement of GNOM activity for MIZ1 function. Plant Physiol 157:1209–1220

OIV (2017) Focus OIV 2017 Distribution of the world’s grapevine varieties. In: OIV (ed). OIV - International organization of vine and wine 18 rue d’Aguesseau F-75008 Paris – France www.oiv.in last accessed Feb 2, 2020.

OIV (2017a) 2017 World Vitiviniculture Situation. OIV Statistical Report on World Vitiviniculture. International Organisation of Vine and Wine

Palumbo F, Vannozzi A, Magon G, Lucchin M, Barcaccia G (2019) Genomics of Flower Identity in Grapevine (*Vitis vinifera* L.). Frontiers in Plant Science 10: 316 https://doi.org/10.3389/fpls.2019.00316

Pelaz S, Ditta GS, Baumann E, Wisman E, Yanofsky MF (2000) B and C floral organ identity functions require SEPALLATA MADS-box genes. Nature 405:200–203

Pertot I, Caffi T, Rossi V, Mugnai L, Hoffmann C, Grando MS, Gary C, Lafond D, Duso C, Thiery D, Mazzoni V, Anfora G (2017) A critical review of plant protection tools for reducing pesticide use on grapevine and new perspectives for the implementation of IPM in viticulture. Crop Protection 97:70–84

Pieri P, Zott K, Gomès E, Hilbert G (2016) Nested effects of berry half, berry and bunch microclimate on biochemical composition in grape

R Core Team (2013) R: A language and environment for statistical computing, Vienna, Austria http://www.r-project.org/index.html

Ranocha P, Dima O, Nagy R, Felten J, Corratge-Faillie C, Novak O, Morreel K, Lacombe B, Martinez Y, Pfrunder S, Jin X, Renou JP, Thibaud JB, Ljung K, Fischer U, Martinoia E, Boerjan W, Goffner D (2013) Arabidopsis WAT1 is a vacuolar auxin transport facilitator required for auxin homoeostasis. Nat Commun 4:2625

Regner F, Stadlbauer A, Eisenheld C, Kaserer H (2000) Genetic Relationships Among Pinots and Related Cultivars. American Journal of Enology and Viticulture 51:7–14

Reid KE, Olsson N, Schlosser J, Peng F, Lund ST (2006) An optimized grapevine RNA isolation procedure and statistical determination of reference genes for real-time RT-PCR during berry development. BMC Plant Biology 6

Richter R, Gabriel D, Rist F, Töpfer R, Zyprian E (2019) Identification of co-located QTLs and genomic regions affecting grapevine cluster architecture. Theoretical and Applied Genetics 132 (4): 1159–1177

Richter R, Rossmann S, Topfer R, Theres K, Zyprian E (2017) Genetic analysis of loose cluster architecture in grapevine. In: Aurand JM (ed) 40th World Congress of Vine and Wine

Rist F, Herzog K, Mack J, Richter R, Steinhage V, Töpfer R (2018) High-Precision Phenotyping of Grape Bunch Architecture Using Fast 3D Sensor and Automation. Sensors 18:763

Rossmann S, Richter R, Sun H, Schneeberger K, Töpfer R, Zyprian E, Theres K (2019) Mutations in the miR396 binding site of the growth-regulating factor gene VvGRF4 modulate inflorescence architecture in grapevine. The Plant Journal, doi: 10.1111/tpj.14588

Ruehl E, Konrad H, Lindner B, Bleser E (2004) Quality criteria and targets for lonal selection in grapevine.. Acta Horticulturae 625:29–33

Sapkota SD, Chen LL, Yang S, Hyma KE, Cadle-Davidson LE, Hwang CF (2019) Quantitative trait locus mapping of downy mildew and botrytis bunch rot resistance in a Vitis aestivalis-derived ‘Norton’-based population. 1248 edn. International Society for Horticultural Science (ISHS), Leuven, Belgium, pp 305–312

Schmittgen TD, Livak KJ (2008) Analyzing real-time PCR data by the comparative C-T method. Nature Protocols 3:1101–1108

Selim M, Legay S, Berkelmann-Löhnertz B, Langen G, Kogel K-H, Evers D (2012) Identification of suitable reference genes for real-time RT-PCR normalization in the grapevine-downy mildew pathosystem. Plant Cell Reports 31:205–216

Shavrukov YN, Dry IB, Thomas MR (2004) Inflorescence and bunch architecture development in *Vitis vinifera* L. Australian Journal of Grape and Wine Research 10:116–124

Smart R, Robinson M (1991) Sunlight into Wine. A Handbook for Winegrape Canopy Management, Adelaide

Smyth Gordon K (2004) Linear Models and Empirical Bayes Methods for Assessing Differential Expression in Microarray Experiments. Statistical Applications in Genetics and Molecular Biology 3:1–25

Suzuki H, Oshita E, Fujimori N, Nakajima Y, Kawagoe Y, Suzuki S (2015) Grape expansins, VvEXPA14 and VvEXPA18 promote cell expansion in transgenic Arabidopsis plant. Plant Cell, Tissue and Organ Culture (PCTOC) 120:1077–1085

Tello J, Aguirrezabal R, Hernaiz S, Larreina B, Montemayor MI, Vaquero E, Ibáñez J (2015) Multicultivar and multivariate study of the natural variation for grapevine bunch compactness. Australian Journal of Grape and Wine Research 21:277–289

Tello J, Ibáñez J (2014) Evaluation of indexes for the quantitative and objective estimation of grapevine bunch compactness. Vitis 53:9–16

Tello J, Ibáñez J (2017) What do we know about grapevine bunch compactness? A state-of-the-art review. Australian Journal of Grape and Wine Research 24:6–23

Tello J, Torres-Perez R, Grimplet J, Ibáñez J (2016) Association analysis of grapevine bunch traits using a comprehensive approach. Theoretical and Applied Genetics 129:227–242

Upadhyay A, Jogaiah S, Maske SR, Kadoo NY, Gupta VS (2015) Expression of stable reference genes and SPINDLY gene in response to gibberellic acid application at different stages of grapevine development. Biologia Plantarum 59:436–444

Vail ME, Marois JJ (1991) Grape cluster architecture and the susceptibility of berries to *Botrytis cinerea*. Phytopathology 81:188–191

Zhang L-Y, Bai M-Y, Wu J, Zhu J-Y, Wang H, Zhang Z, Wang W, Sun Y, Zhao J, Sun X, Yang H, Xu Y, Kim S-H, Fujioka S, Lin W-H, Chong K, Lu T, Wang Z-Y (2009) Antagonistic HLH/bHLH Transcription Factors Mediate Brassinosteroid Regulation of Cell Elongation and Plant Development in Rice and *Arabidopsis*. The Plant Cell 21:3767–3780

