## Supplementary material for "Same same but different: Cluster architecture variation in five ‘Pinot Noir’ clonal selection lines correlates with differential expression of three transcription factors and further growth related genes": Online resource 1

Online resource 1 Weather recording stations, plant protection schedule, average climate conditions and plant vigor at three trial fields

Average air temperature and precipitation were recorded with the nearest weather stations to the trial field region, at comparable latitude level to the trial field during the period April to September. Vegetative vigor was estimated with the weight of the pruned wood per vine.

| Trial field location (Trial field management) | Weather station (Identifier) | latitude over zero: | Trail field established / vine spacing | Plant protection schedule | Season | Average Air temp. [°C] | Average precipitation Apr.-Sept. [mm/m$^2$]¶ | Pruning wood weight [kg] |
|---|---|---|---|---|---|---|---|---|
| Baden<br>48°07'15.9"N<br>7°37'06.0"E<br>(integrated) | König-schaffhausen (84) | 185 m | 1997<br>2.0m*1.1m | BBCH 17-65 sulfur, synthetic fungicides<br>BBCH 65-81 synthetic fungicides every 10-12 days | 2015<br>2016<br>2017 | 12.0<br>11.2<br>11.7 | 47.1<br>67.9<br>53.5 | 1.176$^{a}$<br>1.096$^{a}$<br>- |
| Hesse<br>49°37'28.7"N<br>8°38'54.0"E<br>(integrated) | Hirschberg (135) | 100 m | 1995<br>1.8m*1.0m | BBCH 17-65 sulfur, synthetic fungicides<br>BBCH 65-81 synthetic fungicides every 10-12 days | 2015<br>2016<br>2017 | 12.0<br>11.4<br>11.0 | 50.5<br>64.9<br>73.1 | 0.720$^{b}$<br>0.795$^{b}$<br>- |
| Palatinate<br>49°13'07.8"N<br>8°02'40.5"E<br>(organic) | Siebeldingen (88) | 192 m | 2003<br>2.0m*1.0m | BBCH 17-81 copper, sulfur,<br>BBCH 79-85 copper, carbonates every 7 days | 2015<br>2016<br>2017 | 11.7<br>10.8<br>11.0 | 36.0<br>48.5<br>53.3 | 0.408$^{c}$<br>0.504$^{c}$<br>- |

Letters given in superscript form, indicate significant differences between measurement records according to ANOVA α=0.05
