## Supplementary material for "Same same but different: Cluster architecture variation in five ‘Pinot Noir’ clonal selection lines correlates with differential expression of three transcription factors and further growth related genes": Online resource 2

Online resource 2 Temporal and spatial trial layout of samples used in gene expression experiments.

Rachis samples (three unrelated biological repeats) were taken twice, at pre bloom (phenological stage BBCH57) and at past bloom (BBCH71; see Figure 3) at the three locations Palatinate (P), Hesse (H) and Baden (B) during the seasons 2015-2017. The sampling dates ranged over up to 16 days due to the fixed target for cumulated degree day (CDD) sum (400° BBCH57 and 700° BBCH71) for a phenological stage according to[57]

| Sampling schedule | | | | | |
|---|---|---|---|---|---|
| Location | BBCH | Season | Day of year | Sampling | °CDD start at BBCH09 |
| P | 57 | 2015 | 152 | 01.06.2015 | 421.2 |
| P | 57 | 2016 | 161 | 09.06.2016 | 415.69 |
| P | 57 | 2017 | 154 | 03.06.2017 | 416.65 |
| H | 57 | 2015 | 149 | 29.05.2015 | 398.27 |
| H | 57 | 2016 | 157 | 05.06.2016 | 447.21 |
| H | 57 | 2017 | 152 | 01.06.2017 | 401.87 |
| B | 57 | 2015 | 146 | 31.05.2015 | 390.33 |
| B | 57 | 2016 | 159 | 07.06.2016 | 422.07 |
| B | 57 | 2017 | 150 | 30.05.2017 | 420.15 |
| P | 71 | 2015 | 177 | 26.06.2015 | 714.83 |
| P | 71 | 2016 | 186 | 04.07.2016 | 708.59 |
| P | 71 | 2017 | 177 | 26.06.2017 | 712.11 |
| H | 71 | 2015 | 176 | 25.06.2015 | 725.96 |
| H | 71 | 2016 | 178 | 26.06.2016 | 703.37 |
| H | 71 | 2017 | 171 | 22.06.2017 | 677.38 |
| B | 71 | 2015 | 170 | 19.06.2015 | 698.01 |
| B | 71 | 2016 | 182 | 30.06.2016 | 712.93 |
| B | 71 | 2017 | 171 | 20.06.2017 | 711.26 |
