## Supplementary material for "Same same but different: Cluster architecture variation in five ‘Pinot Noir’ clonal selection lines correlates with differential expression of three transcription factors and further growth related genes": Online resource 3

Online resource 3 Primers for the amplification of candidate genes and reference genes used in this study.

[1]Amplicon length of the product based on the reference genome PN40024 assembly version 12x v2

| Gene ID | Forward sequence 5' -->3' | Reverse sequence 5' -->3' | [1]Amplification product [bp] | Bibliography |
|---|---|---|---|---|
| VIT_00s0313g00070 | AGGTTGAGCAAGGAAGTTGCA | CTCGGCTCAATCCAGCTTCA | 127 | 1 |
| VIT_01s0010g01810 | TCGCCGTTGTCCGAGTTT | ACTTCCACTCCACCACCT | 151 | 2 |
| VIT_01s0010g02430 | CAAGATGAGGGTGTTAAATCGT | ACCTCATTTGTTGCCTTGCT | 119 | 2 |
| VIT_01s0011g06410 | CTCCATGCGGGTCCTTGT | GTGCGTTGGTTTCTGGGATT | 108 | 1 |
| VIT_01s0026g02030 | AAGCCAAAAGCGCAGACA | GCAATAGGCGCTCCGACAA | 118 | 3 |
| VIT_01s0127g00260 | GGCGCGCAAGAAGATCAGAGA | CCCACGCTTGCCAAATAACAT | 199 | 2 |
| VIT_01s0127g00710 | CCCACCTCCTTTATGACCGCTA | CAAGAAAATCCTCCATCAACCGT | 232 | 2 |
| VIT_01s0127g00870 | AGGCGTCTTTGCTTCGGTATT | CGCATTTTGAGCGGCAAGT | 134 | 2 |
| VIT_01s0146g00400 | TCCCACTCCGACACCACCTT | TCTTCCTTGGCTTTCTTGCCGTTT | 131 | 2 |
| VIT_01s0146g00480 | CCGCCATGGAACTTGATTTCT | GCGAACGGCGGATTATTCT | 196 | 2 |
| VIT_02s0012g00990 | GCTGCCACACCTTACTCAT | ATGTACTTACCCCAACAGATGTC | 204 | 2 |
| VIT_02s0012g01380 | GAGACTCCGGCCACCAACAA | GCCCAGCCTTCACCACATTT | 128 | 2 |
| VIT_02s0012g01400 | CCTCGATTCATTCCGCTTCT | CGGCTGCTGATGCTTCTT | 85 | 2 |
| VIT_02s0025g03010 | GGCTGCGAGAGAGTCGTTAAA | ACCTTTTCCATCCCCAGATCCA | 83 | 2 |
| VIT_02s0025g03140 | CCCGGTTTGACATTTCTCAT | CCTCTTGCACTTCGAATCCT | 68 | 2 |
| VIT_02s0025g03180 | CAACATGGTCCCTGCAATC | GGTTGGAGATGGAGCTTCTG | 190 | 2 |
| VIT_02s0025g04340 | CCGAGTGAAATAAGGCATGT | ATAATTGAGGAGGGCTCACA | 41 | 2 |
| VIT_02s0025g04660 | TTGACTGCTGCTCTTGTGCTT | CCACTCCCAAAAACAGAACCTT | 133 | 2 |
| VIT_02s0025g04720 | CCCTGAAGACAAGCGCGATA | GGGTACCATGTTGTGGAGGATGAAG | 298 | 2 |
| VIT_02s0154g00320 | CCTTCTCCTTGCCCTAAACCT | GGTGGCTTTTTGTGGTGGTTTTT | 102 | 2 |
| VIT_02s0154g00380 | CAGCCTCCTCTACAACCT | CTGCTGCTGCTTCTTCTT | 130 | 2 |
| VIT_02s0241g00030 | CTTCAGTCTTCACCTACTGTGA | AGAAGCTTCTTTTGATACCGATAG | 70 | 2 |
| VIT_03s0097g00700 | GGCCTTATGGGGAGAACCTT | TGCCGCAGTGCCTGTAAA | 56 | 2 |
| VIT_04s0008g00180 | CCCTGGACTGTTTCTGTTGCT | AGGACTGCTGGGGGCAAAA | 128 | 2 |
| VIT_04s0008g00370 | CAAGCAAGGAGAGCCAGACA | CCCGTCACAAGCTCAAGCAA | 133 | 2 |
| VIT_04s0008g01100 | CCCCTTGATGGCCAAGTAT | GGAGAGGGGATGCTGAGAT | 197 | 2 |
| VIT_04s0008g01810 | GCTGCAGATTGAGGTGGTT | GTCTGTTCGCCCTGGAAT | 149 | 2 |
| VIT_04s0008g01910 | TCTCCCTCTCCCTCGTCTTC | CATCCTCACCCCCCACTTCA | 210 | 2 |
| VIT_04s0008g02900 | GGGAGGAATTGAAGGCTATGG | GCACCAATGCGCAGCAAA | 162 | 2 |
| VIT_04s0008g02920 | GTGGCTCCCCAGTTAGTGAT | ACCCACCGACAGTTCTTTTG | 145 | 2 |
| VIT_04s0008g04050 | CCTCACACTCCCATGCCCAAA | CCCAAACAAAAAGCAGCAGCAGAA | 89 | 2 |
| VIT_04s0008g04200 | CGAGAGTGCCCAAGAGGTT | CGCATGACCCTGGCAGAA | 108 | 2 |
| VIT_04s0008g05150 | TCTCGCCCAAGGGGTTTT | CTGAAACACTCCATCCTGCTT | 145 | 2 |
| VIT_04s0008g05770 | GGCCGGAAAGGGAGGTTAT | CGCCAGCCGACTTCAAGA | 88 | 2 |
| VIT_04s0008g05830 | GTCTCAGATCGCGTCATTGT | TTGTGGACAGCTCCTGCTT | 93 | 2 |
| VIT_04s0008g06670 | CCCAATCCGATTCTCTCAACAA | CCCTCCTCACCTTCAACAC | 123 | 2 |
| VIT_04s0023g03070 | GGACCTAAGCTGGAACAAG | CACCGTTGCAGGAATCTT | 118 | 1 |
| VIT_04s0069g00790 | ATCCCAGCAAAGACATCAGT | AAACAGAACCAGGCCCAAGA | 212 | 2 |
| VIT_04s0079g00260 | ACCACAAGCCTGCAATTTT | GGCTCTGACCTCAAGGTT | 112 | 2 |
| VIT_07s0031g01850 | TGGGCAGCTAGGAGGAAGATT | GTGGGGGATGCAGTTATGGT | 75 | 1 |
| VIT_08s0007g01310 | ATGGGAAGAGCTGGTTTGG | AGCGGCTAGTGTTCAAATCC | 42 | 2 |
| VIT_08s0007g01320 | GGGGCCGATTCTCAACAGT | ACCACCTCATGGACCTTCCT | 142 | 2 |
| VIT_08s0007g01350 | CTCCTCCTCCTCACGACAGA | CACGCCATCACAGCACTT | 76 | 2 |
| VIT_08s0007g01360 | AACGCCAAACCAGGGACTACA | ACTCTGACTCTCGCCTTCACT | 60 | 2 |
| VIT_08s0007g01370 | GGCGGCCAGCGACAAGA | GGCAGCTTGGGTTCTGGAT | 80 | 2 |
| VIT_08s0040g00040 | GAGGGTCGTCAGGATTTGGA | GCCCTGCACTTACCATCTCTA | 71 | 4 |
| VIT_08s0040g01710 | ACTGGATTTGGTGCGACTT | CGTGTGGCATGAGTCTGTT | 117 | 2 |
| VIT_08s0058g00930 | TCGGACGGGGAAAAGTATGCAA | CCTGGGGCCAACTCTACAAT | 125 | 2 |
| VIT_08s0058g00990 | TGGGTGCTTCTTTGCTTCGT | CGCCCGCATTCTTTTCACT | 111 | 2 |
| VIT_09s0070g00470 | TGCCAAAAGGGACCTCTGAT | TCGGGAGGAGGAAGAGGAGCTA | 113 | 1 |
| VIT_11s0016g03710 | GTCCGAATCGGCTGCTTGAA | TCGGGTTCCATCGCACTT | 88 | 2 |
| VIT_12s0059g00190 | CTCCGGCCAGCTCCAACA | GCCCCTACTCTTGCCCTAAAC | 153 | 5 |
| VIT_14s0066g01060 | CCACCTACAGAACTCCCAAAA | TATCCCTCCCTAGACCTCCAAT | 158 | 2 |
| VIT_14s0066g01390 | ATTTGACTCGGGGAAAGCA | TGGCAGCAAGTGACTGATG | 110 | 2 |
| VIT_14s0083g00410 | CCTTCCCAACCTCCCTTTC | CCTCTCCAACCCCATCATCAC | 173 | 6 |
| VIT_14s0108g00700 | AGTGCGAGTGATGAACAGAGA | GGGCTGCTGCGTATAGTG | 184 | 2 |
| VIT_14s0108g00740 | CGCCATTTCCATGCTTCAC | CAAAACAACACTCGCACACAATC | 139 | 2 |
| VIT_14s0219g00230 | CCGGCTGGTGGACAGTAT | AGAGATGGTTATGGCGGTGGAT | 140 | 7 |
| VIT_15s0048g01750 | CCACCACTCTCTACCAAACC | CCGACCTTGCCACCTTTCA | 85 | 2 |
| *VvGRF4* | ACCAACCAATCCCAATTCCA | TTCGCCTACCTCGGGTTT | 102 | 2 |
| VIT_17s0000g02470 | GGTCCCTGCTTCTCAGTCT | TTGCCTGCGCCTGGTTGTA | 121 | 2 |
| VIT_17s0000g03550 | GGAGAGAGAAAGGCTCGAGTT | AGCATGGAAAGGCGATCAT | 104 | 2 |
| VIT_17s0000g03750 | ACAGAGAGGGGAGAGCTT | TTGTACCACCTGAGATTTGCT | 159 | 2 |

| | | | | |
|---|---|---|---|---|
| VIT_17s0000g04470 | TCCGCCCTGTGTTCTTCT | AACAACTTTCCGATTCCAGATAC | 60 | 2 |
| VIT_17s0000g05000 | GTCGCCTTCCTGCTCAATC | CGGGGCCAAATCCATTGT | 151 | 2 |
| VIT_17s0000g05070 | TCTCTCTCCATAACCTCCCTCAAAC | CCATTAGCGGTGGCAGAAC | 159 | 2 |
| VIT_17s0000g05570 | GCAGGCTTCCCACTTCAAA | CGCTCATCTTGTCCACCAT | 94 | 2 |
| VIT_17s0000g07350 | GAGGATGTGCTGAGGATGGA | TGTGGTCGCATAGCCGTTT | 118 | 2 |
| VIT_17s0000g09190 | AGGGTTCTTGCTGTGGAT | ACACAACTCCCCTAACTTCAC | 66 | 2 |
| VIT_17s0000g09310 | ACCCCCGATGACTACCTTT | CCCTGTGCTTTTGCTGGAT | 169 | 2 |
| VIT_17s0000g09470 | AATTGTCACAGCTTCACCCAAAG | CGCGGTCCACTTGGCTTATC | 164 | 2 |
| VIT_17s0000g09790 | GGGTTGGATGTTTTTGCAAGAT | CCGCCTACTTCGCTTCTTC | 97 | 2 |
| VIT_17s0000g10430 | TTCTCGTTGAGGGCTATT | CCACAGACTTCATCGGTGACA | 70 | 4 |
| VIT_17s0053g00990 | CTTCTATGGCGGGGGTGAT | GCCACAGCTCAACCCATT | 133 | 8 |
| VIT_18s0001g03160 | CGCCTTTCGCACTTGTTC | GGAAGCCAAGCACCATTATTTT | 84 | 2 |
| VIT_18s0001g03540 | GGGCTACCAACATTCTCTACAC | TCCCCAAAAGCCCAATAAACAG | 167 | 2 |
| VIT_18s0001g04890 | TGTGCCGGTGCCTTCTTT | CCTTCTACGCTGGGCCTAA | 118 | 2 |
| VIT_18s0001g04910 | ATCTGCGGCTTGCATTCAC | AGCTCCACCCATAAAACCAACA | 128 | 2 |
| VIT_18s0001g05060 | CAAGCCTCAACTGCTCATAC | CACATCAACACAACCAGTGAAC | 165 | 2 |
| VIT_18s0001g05800 | GGCATTGACTGGGACCAAAA | CCACCTCTTCTGCATCTCT | 140 | 2 |
| VIT_18s0001g07340 | CCCGGTCAGCTTATGTTCAT | AGTGTTGGGGGAGAAGGT | 104 | 2 |
| VIT_18s0001g07460 | GCAGATGAGGGGAGAGGATA | GTGGCGATCTCGGTCATT | 129 | 1 |
| VIT_18s0001g09230 | ACAAGCGATGCCACTACGAA | GGCAGGTTGAGGTCGAAGT | 106 | 2 |
| VIT_18s0001g09400 | CGGATTGCTGGTTCGTCAT | GTCCTTCGTTGCGTCCTT | 116 | 2 |
| VIT_18s0001g09510 | AGGGAGGCAGAAGACGATGA | GTCCCAGCCGAGGTATCTGT | 132 | 2 |
| VIT_18s0001g09910 | CGAAAGAAGCCAACAGCAT | CACCGTTTCTGGCGCATA | 140 | 2 |
| VIT_18s0001g10130 | CACCCGTGAAGGCAAGTC | CGCCGTCTTTGTCATGTT | 83 | 2 |
| VIT_18s0001g10610 | AAACATGCCTCGTCATTGGAA | CGCCGTTTTTGTCATGGT | 119 | 2 |
| VIT_18s0001g10640 | CCGTACGTGCCTAGATTAAAGAA | CCAAGCATCCCCAAATGGAA | 44 | 2 |
| VIT_18s0001g11160 | GTTCGTTTGGGCTGTGTACT | CTCCTCGTCTGACATTTGCTT | 76 | 2 |
| VIT_19s0015g00270 | CGGAGAGTGCTGCTGATGAT | GCTTGACTTTTTCGGGTTTTCGT | 149 | 2 |
| VIT_19s0015g00490 | ACGGAACCGGAGAAGACACT | CCCCATCAGAATCGCCATCT | 108 | 2 |
| VIT_19s0015g01230 | CGTTGTGGAAATAGCTGTGGAT | AATGGGTGGTGGTGGATTG | 87 | 2 |
| VIT_19s0015g01890 | CATTCATCACCCCCGTCTCT | ATTCCCACATTCCCCAAACTCA | 93 | 1 |


1) JIANG, Y., BAO, L., JEONG, S. Y., KIM, S. K., XU, C., LI, X. & ZHANG, Q. 2012. XIAO is involved in the control of organ size by contributing to the regulation of signaling and homeostasis of brassinosteroids and cell cycling in rice. *Plant J,* 70**,** 398-408.

2) ROSSMANN, S., RICHTER, R., SUN, H., SCHNEEBERGER, K., TOPFER, R., ZYPRIAN, E. & THERES, K. 2019. Mutations in the miR396 binding site of the growth-regulating factor gene VvGRF4 modulate inflorescence architecture in grapevine. *Plant J.*

3) ZHANG, L.-Y., BAI, M.-Y., WU, J., ZHU, J.-Y., WANG, H., ZHANG, Z., WANG, W., SUN, Y., ZHAO, J., SUN, X., YANG, H., XU, Y., KIM, S.-H., FUJIOKA, S., LIN, W.-H., CHONG, K., LU, T. & WANG, Z.-Y. 2009. Antagonistic HLH/bHLH Transcription Factors Mediate Brassinosteroid Regulation of Cell Elongation and Plant Development in Rice and Arabidopsis *The Plant Cell,* 21**,** 3767-3780.

4) SELIM, M., LEGAY, S., BERKELMANN-LÖHNERTZ, B., LANGEN, G., KOGEL, K.-H. & EVERS, D. 2012. Identification of suitable reference genes for real-time RT-PCR normalization in the grapevine-downy mildew pathosystem. *Plant Cell Reports,* 31**,** 205-216.

5) DAL SANTO, S., VANNOZZI, A., TORNIELLI, G. B., FASOLI, M., VENTURINI, L., PEZZOTTI, M. & ZENONI, S. 2013. Genome-Wide Analysis of the Expansin Gene Superfamily Reveals Grapevine-Specific Structural and Functional Characteristics. *Plos One,* 8.

6) CORREA, J., MAMANI, M., MUNOZ-ESPINOZA, C., LABORIE, D., MUNOZ, C., PINTO, M. & HINRICHSEN, P. 2014. Heritability and identification of QTLs and underlying candidate genes associated with the architecture of the grapevine cluster (Vitis vinifera L.). *Theoretical and Applied Genetics,* 127**,** 1143-1162.

7) VARGAS, A. M., FAJARDO, C., BORREGO, J., DE ANDRES, M. T. & IBANEZ, J. 2013. Polymorphisms in VvPel associate with variation in berry texture and bunch size in the grapevine. *Australian* Journal of Grape and Wine Research, 19, 193-207.

8)HOFFMANN, P. 2015. Lockerbeerigkeit bei Klonen von Spätburgunder (Pinot noir) : Analyse von molekularen Markern und der Einfluss von Gibberellin auf die Traubenmorphologie. Kommunikations-, Informations- und Medienzentrum der Universität Hohenheim. http://opus.uni-hohenheim.de/volltexte/2015/1022/
