## Supplementary material for "Same same but different: Cluster architecture variation in five ‘Pinot Noir’ clonal selection lines correlates with differential expression of three transcription factors and further growth related genes": Online resource 4

Online resource 4a Effects of trial location and growing season on important cluster architecture sub traits and compactness indices for the 'Pinot Noir' clones Gm20-13 and FkCH, which were the two reference clones that were sampled across all seasons and locations. Mean and 95% confidence interval were estimated with generalized linear models (n = 120).

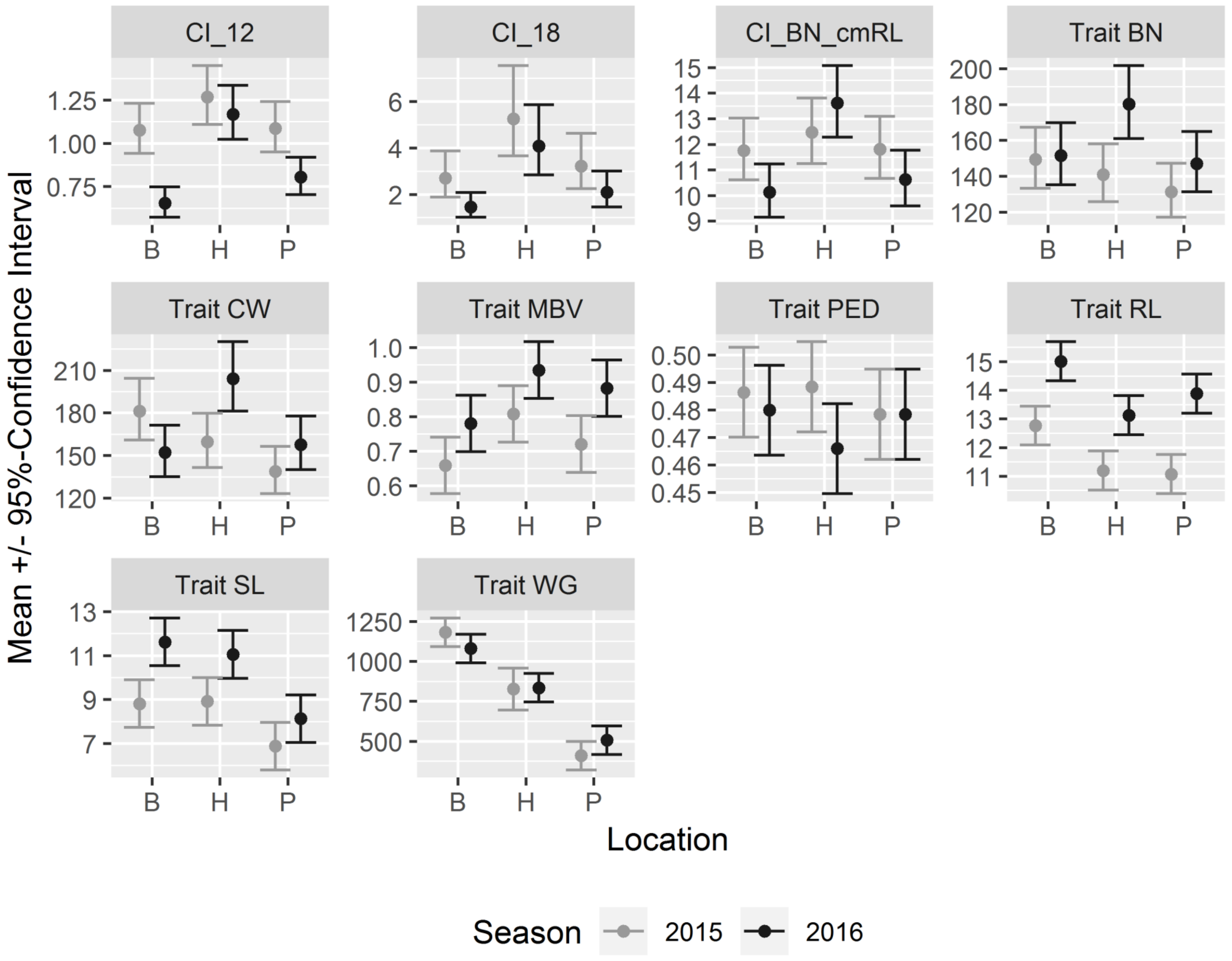


Estimated marginal means for measurements from 2015 and 2016 at three trial fields located in German wine growing regions. Hesse (H) and Palatinate (P) belong to viticulture area A (cool climate). Baden (B) belongs to viticulture area B (moderate climate). For trait abbreviations see table 2.

### Online resource 4b ANOVA results for important cluster architecture sub traits and compactness indices for 'Pinot Noir' clones that were sampled across all seasons and locations. Mean and 95% confidence interval were estimated with generalized linear models.

ANOVA results of a reciprocal design (n = 120) for the measurement records of the 'Pinot Noir' clones Gm20-13 and FkCH at all locations and seasons.
P-values for the effects of trial field location and growing season on cluster architecture sub-traits and compactness indices, obtained from generalized linear models (GLM) with negative binomial (NB) or gamma distribution or ordinary least squares models (OLS) and ANOVA sums of squares type 3 test.
For trait abbreviations see table 2.

| Trait / Index | Model | Clone | Location | Season | Location: Season |
|---|---|---|---|---|---|
| BN | NB GLM | 0.000 | 0.059 | 0.008 | 0.130 |
| CW | Gamma GLM | 0.000 | 0.004 | 0.178 | 0.002 |
| MBV | OLS | 0.000 | 0.002 | 0.000 | 0.868 |
| TBV | NB GLM | 0.000 | 0.000 | 0.000 | 0.195 |
| RD | OLS | 0.004 | 0.000 | 0.133 | 0.151 |
| RL | OLS | 0.019 | 0.000 | 0.000 | 0.436 |
| RW | OLS | 0.000 | 0.006 | 0.000 | 0.367 |
| SL | OLS | 0.722 | 0.000 | 0.000 | 0.362 |
| PL | OLS | 0.125 | 0.208 | 0.081 | 0.101 |
| PED | OLS | 0.882 | 0.746 | 0.155 | 0.378 |
| L1I | OLS | 0.694 | 0.168 | 0.199 | 0.568 |
| L2I | OLS | 0.179 | 0.274 | 0.089 | 0.337 |
| BN_cmRL | Gamma GLM | 0.000 | 0.000 | 0.188 | 0.052 |
| CI_12 | Gamma GLM | 0.000 | 0.000 | 0.000 | 0.000 |
| CI_18 | Gamma GLM | 0.081 | 0.000 | 0.004 | 0.606 |
| WG | Gamma GLM | 0.024 | 0.000 | 0.293 | 0.043 |

ANOVA results for measurement records of twelve 'Pinot Noir' clones (n = 400).

P-values for the effects of trial field location and growing season on cluster architecture sub-traits and compactness indices, obtained from generalized linear models (GLM) with negative binomial (NB) or gamma distribution or ordinary least squares models (OLS) and ANOVA sums of squares type 3 test.
For trait abbreviations see table 2.

| Trait / Index | Model | Clone | Location | Season | Location: Season |
|---|---|---|---|---|---|
| BN | NB GLM | 0.056 | 0.000 | 0.004 | 0.001 |
| CW | Gamma GLM | 0.000 | 0.000 | 0.000 | 0.000 |
| MBV | OLS | 0.000 | 0.009 | 0.000 | 0.000 |
| TBV | NB GLM | 0.000 | 0.000 | 0.009 | 0.327 |
| RD | OLS | 0.000 | 0.000 | 0.009 | 0.327 |
| RL | OLS | 0.000 | 0.000 | 0.000 | 0.244 |
| RW | OLS | 0.000 | 0.000 | 0.000 | 0.007 |
| SL | OLS | 0.000 | 0.004 | 0.000 | 0.527 |
| PL | OLS | 0.000 | 0.008 | 0.062 | 0.107 |
| PED | OLS | 0.000 | 0.662 | 0.000 | 0.368 |
| L1I | OLS | 0.053 | 0.864 | 0.235 | 0.216 |
| L2I | OLS | 0.083 | 0.159 | 0.225 | 0.983 |
| BN_cmRL | Gamma GLM | 0.000 | 0.000 | 0.043 | 0.041 |
| CI_12 | Gamma GLM | 0.000 | 0.000 | 0.001 | 0.000 |
| CI_18 | Gamma GLM | 0.000 | 0.000 | 0.021 | 0.058 |
| WG | Gamma GLM | 0.000 | 0.000 | 0.007 | 0.042 |
