## Supplementary material for "Same same but different: Cluster architecture variation in five ‘Pinot Noir’ clonal selection lines correlates with differential expression of three transcription factors and further growth related genes": Online resource 5

**Online resource 5** Detailed Relative gene expression log $_{(2)}$ FC (-ΔΔt) at BBCH57 and BBCH71 as predicted with a linear model: log $_{(2)}$ FC~ clone*season*location for 'Pinot Noir' clones with divergent cluster architecture. Cluster architecture types were coloured as follows: loosely (LCC) = blue; compact (CCC) = orange; mixed berried (MBC)= green. Trial locations are indicated by capital letters: B = Baden; H = Hesse; P = Palatinate. Season is indicates as YY. Abbreviations for clone names see table 1. The contrast is to clone Gm20-13 (small berries and short rachis features) and the mean of two internal control genes. Genes highlighted in grey show a stable differentially expression between clones with different cluster types over multiple seasons and locations.

**Time point BBCH57**

| featureNames | Fr12L.B.16 | Fr12L.B.17 | Fr12L.H.16 | Fr12L.H.17 | Fr13L.B.16 | Fr13L.B.17 | Fr13L.H.16 | Fr13L.H.17 | Gm1_86.H.15 |
|---|---|---|---|---|---|---|---|---|---|
| VIT_00s0313g00070 | -0,031656349 | -0,107036679 | 0,54639556 | 0,231384978 | 0,08865564 | -0,020899059 | 0,553013992 | 0,154924527 | 0,38177866 |
| VIT_01s0010g01810 | -0,964469159 | -0,764932184 | 0,027643448 | -0,611036565 | 0,217934954 | 0,09250939 | 0,744741132 | 0,08386497 | 1,851029226 |
| VIT_01s0010g02430 | -0,656225297 | -0,002462276 | 1,23545609 | -0,592192924 | -0,561761115 | 0,023213996 | 0,502847536 | -0,592286591 | 2,246717893 |
| VIT_01s0011g06410 | -0,006727315 | 0,052504744 | 0,416454565 | 0,153626523 | -0,111185491 | -0,086447017 | 0,566317066 | 0,03126465 | 0,91977041 |
| VIT_01s0026g02030 | 0,272444298 | 0,098405988 | 0,326007755 | 0,104850405 | 0,220005487 | 0,357543924 | -0,290973388 | 0,230318739 | 0,699898026 |
| VIT_01s0127g00260 | -0,118566782 | -0,142267052 | 1,35373321 | 1,02111434 | -0,338604421 | 0,101177505 | 1,259894094 | 0,414097064 | 1,353142423 |
| VIT_01s0127g00710 | 11,54505947 | 12,01915593 | -0,410288356 | -0,109862239 | 11,2290477 | 11,86499069 | -0,476176356 | 0,296507992 | -2,210214996 |
| VIT_01s0127g00870 | 0,070045928 | 0,395025198 | 2,222709943 | -0,012553765 | 0,154200554 | 0,432704994 | 2,411442624 | -0,127833196 | 3,446658558 |
| VIT_01s0146g00400 | -0,550945055 | 0,382257203 | 0,249830271 | 0,106817861 | -0,407457496 | 0,29108886 | 0,375914461 | -0,185197893 | 2,079498708 |
| VIT_01s0146g00480 | 0,310834685 | 0,197563723 | 0,453616261 | -0,406441492 | -0,858364043 | -0,27396214 | 1,140483182 | -0,55764025 | 0,627068381 |
| VIT_02s0012g00990 | 11,74472123 | 6,591205447 | -0,574485744 | -0,250968234 | 11,37690063 | 6,363908187 | -0,219275483 | 0,186923144 | 0,438548194 |
| VIT_02s0012g01380 | -0,139707379 | -0,065609045 | -0,399206084 | 0,353829958 | 0,068518052 | 0,005273297 | -0,919859096 | [illegible] | [illegible] |
| VIT_02s0025g03180 | -0,20745548 | 0,169105872 | 0,133570743 | 0,070112285 | -0,326524738 | 0,019882194 | 0,301013446 | [illegible] | [illegible] |
| VIT_02s0025g04340 | 0,236135263 | 0,620549441 | -0,596499771 | -0,267172295 | 0,591705922 | 0,633752116 | 0,206377609 | [illegible] | [illegible] |
| VIT_02s0025g04660 | 0,157958373 | -0,326724378 | -0,276270022 | 0,115540083 | 0,343029899 | -0,187630268 | 0,015212426 | [illegible] | [illegible] |
| VIT_02s0025g04720 | -0,302612052 | 7,049005733 | 1,251443472 | 0,525833246 | -0,571810383 | 6,97142884 | 1,071737771 | [illegible] | [illegible] |
| VIT_02s0154g00320 | 0,085751446 | 0,308233924 | 0,33809262 | 0,045958565 | -0,17596708 | 0,345096758 | 0,895482171 | [illegible] | [illegible] |
| VIT_02s0154g00380 | 0,897352703 | -0,634235376 | -0,167701377 | 0,100350105 | 0,20809175 | -0,506690075 | 0,009593439 | [illegible] | [illegible] |
| VIT_02s0241g00030 | -0,251226565 | 0,068386142 | 1,004582935 | 0,067178805 | 0,096909092 | 0,206093229 | 0,699289767 | [illegible] | [illegible] |
| VIT_03s0097g00700 | 0,121585006 | 0,128173448 | -0,494576584 | 0,049623118 | -0,049364495 | 1,608845057 | 0,087926571 | [illegible] | [illegible] |
| VIT_04s0008g00180 | -0,479884152 | 0,34725298 | 0,59322401 | 0,422324408 | -0,105154598 | 0,272443024 | 0,642264587 | [illegible] | [illegible] |
| VIT_04s0008g00370 | 0,477350296 | 0,298829789 | -0,061137227 | 0,567286456 | 0,34129501 | 0,35854172 | -0,155878668 | [illegible] | [illegible] |

[illegible]

**Time point BBCH71**

[illegible]
