## Supplementary material for "Same same but different: Cluster architecture variation in five ‘Pinot Noir’ clonal selection lines correlates with differential expression of three transcription factors and further growth related genes": Online resource 6

Online resource 6 Relative expression of *VvGRF4* a,c) in 'Pinot Noir' clones with different cluster architecture b,d) in clones grouped according to their cluster architecture type: loose (LCC) mixed berried (MBC) and compact (CCC). The gene expression was measured as $\log_{(2)}$ fold change at BBCH57 (a,b) and BBCH71 (c,d) in contrast to the 'Pinot Noir' clone Gm20-13 with small berries and short rachis.

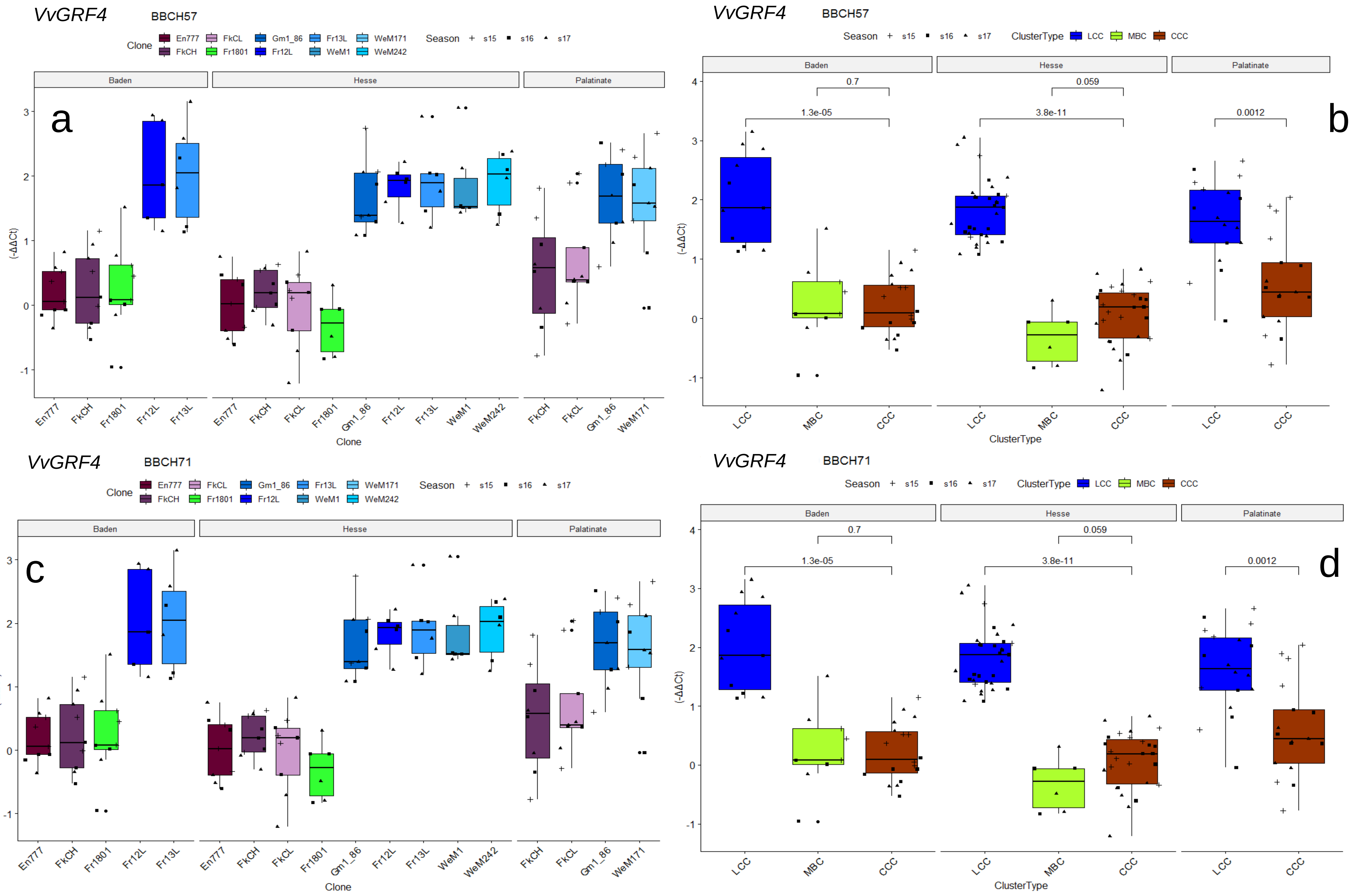
