## Supplementary material for "Same same but different: Cluster architecture variation in five ‘Pinot Noir’ clonal selection lines correlates with differential expression of three transcription factors and further growth related genes": Online resource 7

Online resource 7 Coefficients of Spearman correlation between the relative expression of selected genes and key sub traits of cluster architecture, vegetative vigor measured as wood gain (WG) and the coefficients of correlation between two given gens. The gene expression relative to GAPDH and UBIc as $\log_{(2)}$ of the fold change was measured just before flowering (BBCH57) and just after flowering (BBCH71). The measurement results for cluster architecture sub traits of 'Pinot Noir' clones were recorded at ripe grape clusters (BBCH89). The wood gain was recorded after leaves had fallen (BBCH97). Spearman correlation (r) is significant with *p <0.05, **p <0.01, ***p <0.001 and ****p <0.0001. For trait abbreviations see Table 2. For gene functions see online resource 4. Significant and coherently directed correlations over two seasons are labeled in color. For information regarding the expression clusters c1 to c5 see Figure 2. Candidate genes with putative transcription factor function are labeled in bold.

| Sampling | Season | Gene ID | BN | CW | MBV | PED | RL | SL | WG | VIT_04s0008g01100 c1 | ***VvGRF4*** c2 | VIT_18s0001g03160 c1 | gene expression cluster |
|---|---|---|---|---|---|---|---|---|---|---|---|---|---|
| BBCH57 | 2015 | VIT_04s0008g01100 | 0.72* | 0.60 | -0.94**** | -0.82** | -0.49 | -0.10 | 0.50 | | -0.83** | 0.79** | c1 |
| BBCH57 | 2016 | VIT_04s0008g01100 | 0.40 | -0.48 | -0.78** | -0.93*** | -0.59 | 0.31 | 0.77** | | -0.90*** | 0.95**** | c1 |
| BBCH57 | 2015 | ***VvGRF4*** | -0.37 | -0.21 | 0.87** | 0.92*** | 0.50 | -0.07 | -0.78** | -0.83** | | -0.98**** | c2 |
| BBCH57 | 2016 | ***VvGRF4*** | -0.33 | 0.67* | 0.90*** | 0.89*** | 0.41 | -0.56 | -0.93*** | -0.90*** | | -0.95**** | c2 |
| BBCH57 | 2015 | VIT_18s0001g03160 | 0.37 | 0.21 | -0.83** | -0.83** | -0.38 | 0.16 | 0.83** | 0.79** | -0.98**** | | c1 |
| BBCH57 | 2016 | VIT_18s0001g03160 | 0.31 | -0.64* | -0.88*** | -0.84** | -0.45 | 0.42 | 0.88*** | 0.95**** | -0.95**** | | c1 |

| Sampling | Season | Gene ID | BN | CW | MBV | PED | RL | SL | WG | VIT_01s0010g02430 c5 | **VIT_01s0026g02030** c4 | VIT_01s0127g00870 c5 | VIT_02s0025g04720 c5 | VIT_04s0008g01100 c1 | VIT_08s0007g01370 c1 | ***VvGRF4*** c2 | VIT_17s0000g03750 c5 | **VIT_17s0000g05000** c3 | VIT_17s0053g00990 c5 | VIT_18s0001g03160 c1 | VIT_18s0001g03540 c3 | VIT_18s0001g04890 c1 | VIT_18s0001g05060 c5 | VIT_18s0001g11160 c3 | gene expression cluster |
|---|---|---|---|---|---|---|---|---|---|---|---|---|---|---|---|---|---|---|---|---|---|---|---|---|---|
| BBCH71 | 2015 | VIT_01s0010g02430 | -0.75**** | -0.68*** | 0.90**** | 0.63** | -0.45* | -0.81**** | -0.97**** | | 0.92**** | 0.99**** | 0.95**** | -0.97**** | -0.93**** | 0.93**** | 0.93**** | 0.70*** | 0.95**** | -0.97**** | -0.45* | -0.99**** | 0.99**** | 0.98**** | c5 |
| BBCH71 | 2016 | VIT_01s0010g02430 | 0.61** | 0.70*** | 0.82**** | 0.63** | 0.16 | -0.62** | -0.54* | | 0.86**** | 0.96**** | 0.92**** | -0.74**** | -0.87**** | 0.97**** | 0.83**** | 0.72*** | 0.95**** | -0.88**** | -0.57** | -0.73*** | 0.92**** | 0.82**** | c5 |
| BBCH71 | 2015 | **VIT_01s0026g02030** | -0.70*** | -0.60** | 0.85**** | 0.72*** | -0.24 | -0.71*** | -0.89**** | 0.92**** | | 0.92**** | 0.88**** | -0.90**** | -0.90**** | 0.97**** | 0.89**** | 0.79**** | 0.90**** | -0.89**** | -0.25 | -0.91**** | 0.90**** | 0.92**** | c4 |
| BBCH71 | 2016 | **VIT_01s0026g02030** | 0.81**** | 0.87**** | 0.77**** | 0.48* | -0.01 | -0.52* | -0.61** | 0.86**** | | 0.82**** | 0.98**** | -0.67*** | -0.72*** | 0.87**** | 0.89**** | 0.89**** | 0.86**** | -0.91**** | -0.56** | -0.62** | 0.97**** | 0.89**** | c4 |
| BBCH71 | 2015 | VIT_01s0127g00870 | -0.71*** | -0.64** | 0.88**** | 0.65** | -0.44* | -0.81**** | -0.96**** | 0.99**** | 0.92**** | | 0.96**** | -0.96**** | -0.91**** | 0.95**** | 0.95**** | 0.73*** | 0.95**** | -0.98**** | -0.45* | -0.98**** | 0.99**** | 0.97**** | c5 |
| BBCH71 | 2016 | VIT_01s0127g00870 | 0.52* | 0.69*** | 0.92**** | 0.74**** | 0.23 | -0.69*** | -0.70*** | 0.96**** | 0.82**** | | 0.87**** | -0.86**** | -0.89**** | 0.96**** | 0.71*** | 0.66*** | 0.97**** | -0.92**** | -0.74**** | -0.85**** | 0.88**** | 0.73*** | c5 |
| BBCH71 | 2015 | VIT_02s0025g04720 | -0.62** | -0.54* | 0.81**** | 0.61** | -0.44* | -0.80**** | -0.94**** | 0.95**** | 0.88**** | 0.96**** | | -0.93**** | -0.88**** | 0.92**** | 0.98**** | 0.79**** | 0.98**** | -0.99**** | -0.47* | -0.95**** | 0.97**** | 0.93**** | c5 |
| BBCH71 | 2016 | VIT_02s0025g04720 | 0.77**** | 0.81**** | 0.76**** | 0.51* | 0.00 | -0.57** | -0.59** | 0.92**** | 0.98**** | 0.87**** | | -0.71*** | -0.73*** | 0.92**** | 0.91**** | 0.83**** | 0.91**** | -0.93**** | -0.57** | -0.64** | 0.98**** | 0.87**** | c5 |
| BBCH71 | 2015 | VIT_04s0008g01100 | 0.77**** | 0.68*** | -0.87**** | -0.66*** | 0.39 | 0.73*** | 0.94**** | -0.97**** | -0.90**** | -0.96**** | -0.93**** | | 0.98**** | -0.87**** | -0.89**** | -0.60** | -0.95**** | 0.95**** | 0.34 | 0.98**** | -0.97**** | -0.95**** | c1 |
| BBCH71 | 2016 | VIT_04s0008g01100 | -0.31 | -0.56** | -0.88**** | -0.79**** | -0.21 | 0.75**** | 0.87**** | -0.74**** | -0.67*** | -0.86**** | -0.71*** | | 0.70*** | -0.74**** | -0.45* | -0.42* | -0.83**** | 0.88**** | 0.92**** | 0.97**** | -0.74**** | -0.46* | c1 |
| BBCH71 | 2015 | VIT_08s0007g01370 | 0.77**** | 0.66*** | -0.86**** | -0.69*** | 0.30 | 0.67*** | 0.91**** | -0.93**** | -0.90**** | -0.91**** | -0.88**** | 0.98**** | | -0.86**** | -0.83**** | -0.56** | -0.92**** | 0.91**** | 0.21 | 0.93**** | -0.92**** | -0.92**** | c1 |
| BBCH71 | 2016 | VIT_08s0007g01370 | -0.44* | -0.64** | -0.88**** | -0.70*** | -0.31 | 0.55** | 0.53* | -0.87**** | -0.72*** | -0.89**** | -0.73*** | 0.70*** | | -0.88**** | -0.63** | -0.64** | -0.88**** | 0.73*** | 0.54** | 0.76**** | -0.73*** | -0.73*** | c1 |
| BBCH71 | 2015 | ***VvGRF4*** | -0.64** | -0.55** | 0.83**** | 0.72*** | -0.31 | -0.76**** | -0.90**** | 0.93**** | 0.97**** | 0.95**** | 0.92**** | -0.87**** | -0.86**** | | 0.94**** | 0.85**** | 0.92**** | -0.92**** | -0.35 | -0.91**** | 0.92**** | 0.92**** | c2 |
| BBCH71 | 2016 | ***VvGRF4*** | 0.62** | 0.72*** | 0.84**** | 0.66*** | 0.18 | -0.58** | -0.55** | 0.97**** | 0.87**** | 0.96**** | 0.92**** | -0.74**** | -0.88**** | | 0.83**** | 0.74**** | 0.97**** | -0.89**** | -0.57** | -0.72*** | 0.89**** | 0.86**** | c2 |
| BBCH71 | 2015 | VIT_17s0000g03750 | -0.58** | -0.49* | 0.78**** | 0.70*** | -0.37 | -0.76**** | -0.90**** | 0.93**** | 0.89**** | 0.95**** | 0.98**** | -0.89**** | -0.83**** | 0.94**** | | 0.81**** | 0.96**** | -0.96**** | -0.42 | -0.92**** | 0.94**** | 0.90**** | c5 |
| BBCH71 | 2016 | VIT_17s0000g03750 | 0.77**** | 0.73*** | 0.56** | 0.24 | -0.17 | -0.44* | -0.30 | 0.83**** | 0.89**** | 0.71*** | 0.91**** | -0.45* | -0.63** | 0.83**** | | 0.84**** | 0.76**** | -0.76**** | -0.27 | -0.37 | 0.88**** | 0.91**** | c5 |
| BBCH71 | 2015 | **VIT_17s0000g05000** | -0.32 | -0.22 | 0.59** | 0.48* | -0.22 | -0.69*** | -0.71*** | 0.70*** | 0.79**** | 0.73*** | 0.79**** | -0.60** | -0.56** | 0.85**** | 0.81**** | | 0.75**** | -0.74**** | -0.39 | -0.68*** | 0.71*** | 0.69*** | c3 |
| BBCH71 | 2016 | **VIT_17s0000g05000** | 0.85**** | 0.88**** | 0.63** | 0.23 | -0.10 | -0.38 | -0.48* | 0.72*** | 0.89**** | 0.66*** | 0.83**** | -0.42* | -0.64** | 0.74**** | 0.84**** | | 0.68*** | -0.74**** | -0.38 | -0.42 | 0.81**** | 0.89**** | c3 |
| BBCH71 | 2015 | VIT_17s0053g00990 | -0.63** | -0.54** | 0.81**** | 0.65*** | -0.40 | -0.77**** | -0.93**** | 0.95**** | 0.90**** | 0.95**** | 0.98**** | -0.95**** | -0.92**** | 0.92**** | 0.96**** | 0.75**** | | -0.98**** | -0.40 | -0.95**** | 0.96**** | 0.92**** | c5 |
| BBCH71 | 2016 | VIT_17s0053g00990 | 0.54** | 0.68*** | 0.88**** | 0.70*** | 0.16 | -0.66*** | -0.65*** | 0.95**** | 0.86**** | 0.97**** | 0.91**** | -0.83**** | -0.88**** | 0.97**** | 0.76**** | 0.68*** | | -0.92**** | -0.67*** | -0.80**** | 0.90**** | 0.79**** | c5 |
| BBCH71 | 2015 | VIT_18s0001g03160 | 0.65** | 0.58** | -0.82**** | -0.61** | 0.46* | 0.81**** | 0.96**** | -0.97**** | -0.89**** | -0.98**** | -0.99**** | 0.95**** | 0.91**** | -0.92**** | -0.96**** | -0.74**** | -0.98**** | | 0.48* | 0.97**** | -0.99**** | -0.94**** | c1 |
| BBCH71 | 2016 | VIT_18s0001g03160 | -0.64** | -0.81**** | -0.89**** | -0.61** | -0.06 | 0.70*** | 0.80**** | -0.88**** | -0.91**** | -0.92**** | -0.93**** | 0.88**** | 0.73*** | -0.89**** | -0.76**** | -0.74**** | -0.92**** | | 0.79**** | 0.80**** | -0.93**** | -0.75**** | c1 |
| BBCH71 | 2015 | VIT_18s0001g03540 | 0.21 | 0.40 | -0.28 | 0.26 | 0.88**** | 0.78**** | 0.51* | -0.45* | -0.25 | -0.45* | -0.47* | 0.34 | 0.21 | -0.35 | -0.42 | -0.39 | -0.40 | 0.48* | | 0.45* | -0.47* | -0.43* | c3 |
| BBCH71 | 2016 | VIT_18s0001g03540 | -0.26 | -0.52* | -0.79**** | -0.65*** | -0.10 | 0.75**** | 0.96**** | -0.57** | -0.56** | -0.74**** | -0.57** | 0.92**** | 0.54** | -0.57** | -0.27 | -0.38 | -0.67*** | 0.79**** | | 0.92**** | -0.62** | -0.29 | c3 |
| BBCH71 | 2015 | VIT_18s0001g04890 | 0.76**** | 0.69*** | -0.90**** | -0.61** | 0.46* | 0.80**** | 0.98**** | -0.99**** | -0.91**** | -0.98**** | -0.95**** | 0.98**** | 0.93**** | -0.91**** | -0.92**** | -0.68*** | -0.95**** | 0.97**** | 0.45* | | -0.99**** | -0.98**** | c1 |
| BBCH71 | 2016 | VIT_18s0001g04890 | -0.27 | -0.54** | -0.88**** | -0.82**** | -0.30 | 0.72*** | 0.86**** | -0.73*** | -0.62** | -0.85**** | -0.64** | 0.97**** | 0.76**** | -0.72*** | -0.37 | -0.42 | -0.80**** | 0.80**** | 0.92**** | | -0.68*** | -0.42 | c1 |
| BBCH71 | 2015 | VIT_18s0001g05060 | -0.72*** | -0.64** | 0.88**** | 0.61** | -0.47* | -0.81**** | -0.98**** | 0.99**** | 0.90**** | 0.99**** | 0.97**** | -0.97**** | -0.92**** | 0.92**** | 0.94**** | 0.71*** | 0.96**** | -0.99**** | -0.47* | -0.99**** | | 0.97**** | c5 |
| BBCH71 | 2016 | VIT_18s0001g05060 | 0.75**** | 0.80**** | 0.76**** | 0.51* | -0.02 | -0.61** | -0.63** | 0.92**** | 0.97**** | 0.88**** | 0.98**** | -0.74**** | -0.73*** | 0.89**** | 0.88**** | 0.81**** | 0.90**** | -0.93**** | -0.62** | -0.68*** | | 0.84**** | c5 |
| BBCH71 | 2015 | VIT_18s0001g11160 | -0.77**** | -0.70*** | 0.92**** | 0.63** | -0.43* | -0.79**** | -0.98**** | 0.98**** | 0.92**** | 0.97**** | 0.93**** | -0.95**** | -0.92**** | 0.92**** | 0.90**** | 0.69*** | 0.92**** | -0.94**** | -0.43* | -0.98**** | 0.97**** | | c3 |
| BBCH71 | 2016 | VIT_18s0001g11160 | 0.75**** | 0.76**** | 0.66*** | 0.33 | -0.02 | -0.39 | -0.35 | 0.82**** | 0.89**** | 0.73*** | 0.87**** | -0.46* | -0.73*** | 0.86**** | 0.91**** | 0.89**** | 0.79**** | -0.75**** | -0.29 | -0.42 | 0.84**** | | c3 |
