## Supplementary material for "Same same but different: Cluster architecture variation in five ‘Pinot Noir’ clonal selection lines correlates with differential expression of three transcription factors and further growth related genes": Online resource 8

Online resource 8 Variance partition analysis using experimental, biological and technical factors to reveal their fractions of explained variance in relative candidate gene expression log(2) ($\Delta C_t$).
Factors of variance are: cluster type (loose, mixed berried, compact), bio replicates, (biological variance), season, batch (technical variance), location, gene pool (selection background) and clone (11 'Pinot Noir' clones). [1]Median of the fraction of variance explained by an individual factor.

**BBCH57**

| Gene ID | Batch | bio_Replicates | Clone | Cluster_Type | Clone_Pool | Location | Season | Residuals |
|---|---|---|---|---|---|---|---|---|
| VIT_04s0008g01100 | 0.170 | 0.178 | 0.000 | 0.154 | 0.018 | 0.215 | 0.042 | 0.223 |
| *VvGRF4* | 0.008 | 0.083 | 0.017 | 0.584 | 0.000 | 0.002 | 0.181 | 0.125 |
| VIT_18s0001g03160 | 0.000 | 0.044 | 0.011 | 0.134 | 0.000 | 0.163 | 0.260 | 0.388 |
| [1]Median | 0.008 | 0.083 | 0.011 | 0.154 | 0.000 | 0.163 | 0.181 | 0.223 |

**BBCH71**

| Gene ID | Batch | bio_Replicates | Clone | Cluster_Type | Clone_Pool | Location | Season | Residuals |
|---|---|---|---|---|---|---|---|---|
| VIT_01s0010g02430 | 0.014 | 0.317 | 0.000 | 0.229 | 0.021 | 0.013 | 0.096 | 0.310 |
| VIT_01s0026g02030 | 0.012 | 0.329 | 0.000 | 0.325 | 0.000 | 0.014 | 0.010 | 0.310 |
| VIT_01s0127g00870 | 0.015 | 0.282 | 0.000 | 0.423 | 0.000 | 0.010 | 0.100 | 0.169 |
| VIT_02s0025g04720 | 0.134 | 0.159 | 0.000 | 0.264 | 0.000 | 0.011 | 0.069 | 0.363 |
| VIT_04s0008g01100 | 0.049 | 0.136 | 0.000 | 0.272 | 0.000 | 0.061 | 0.167 | 0.315 |
| VIT_08s0007g01370 | 0.000 | 0.079 | 0.000 | 0.279 | 0.000 | 0.000 | 0.243 | 0.399 |
| *VvGRF4* | 0.008 | 0.067 | 0.000 | 0.835 | 0.000 | 0.000 | 0.026 | 0.064 |
| VIT_17s0000g03750 | 0.060 | 0.196 | 0.000 | 0.255 | 0.104 | 0.059 | 0.019 | 0.307 |
| VIT_17s0000g05000 | 0.035 | 0.000 | 0.000 | 0.271 | 0.030 | 0.061 | 0.262 | 0.342 |
| VIT_17s0053g00990 | 0.028 | 0.198 | 0.000 | 0.318 | 0.000 | 0.016 | 0.270 | 0.170 |
| VIT_18s0001g03160 | 0.151 | 0.120 | 0.000 | 0.365 | 0.018 | 0.082 | 0.000 | 0.263 |
| VIT_18s0001g03540 | 0.005 | 0.070 | 0.000 | 0.136 | 0.054 | 0.220 | 0.384 | 0.132 |
| VIT_18s0001g04890 | 0.340 | 0.142 | 0.000 | 0.229 | 0.007 | 0.000 | 0.128 | 0.154 |
| VIT_18s0001g05060 | 0.126 | 0.213 | 0.000 | 0.232 | 0.000 | 0.070 | 0.160 | 0.200 |
| VIT_18s0001g11160 | 0.000 | 0.287 | 0.000 | 0.331 | 0.018 | 0.000 | 0.112 | 0.252 |
| Median | 0.028 | 0.159 | 0.000 | 0.272 | 0.000 | 0.014 | 0.112 | 0.263 |
